# Screening of Relevant Metabolism-Disrupting Chemicals on Pancreatic β-Cells: Evaluation of Murine and Human in Vitro Models

**DOI:** 10.1101/2022.03.22.485270

**Authors:** Ruba Al-Abdulla, Hilda Ferrero, Sergi Soriano, Talía Boronat-Belda, Paloma Alonso-Magdalena

## Abstract

Endocrine-disrupting chemicals (EDCs) are chemical substances that can interfere with the normal function of the endocrine system. EDCs are ubiquitous and can be found in a variety of consumer products such as food packaging materials, personal care and household products, plastic additives, and flame retardants. Over the last decade, the impact of EDCs on human health has been widely acknowledged as they have been associated with different endocrine diseases. Among them, a subset called metabolism-disrupting chemicals (MDCs) are able to promote metabolic changes that can lead to the development of metabolic disorders such as diabetes, obesity, hepatic steatosis, and metabolic syndrome, among others. Despite this, today, there are still no definitive and standardized in vitro tools to support the metabolic risk assessment of existing and emerging MDCs for regulatory purposes. Here, we evaluated two different pancreatic cell-based in vitro systems, the murine pancreatic β-cell line MIN6 as well as the human pancreatic β-cell line EndoC- βH1. Both were challenged with a range of relevant concentrations of seven well-known EDCs (bisphenol-A (BPA), bisphenol-S (BPS), bisphenol-F (BPF), perfluorooctanesulfonic acid (PFOS), di(2-ethylhexyl) phthalate (DEHP), cadmium chloride (CdCl_2_) and dichlorodiphenyldichloroethylene (DDE)). The screening revealed that most of the tested chemicals have detectable deleterious effects on glucose-stimulated insulin release, insulin content, electrical activity, gene expression, and/or viability. Our data provide new molecular information on the direct effects of the selected chemicals on key aspects of pancreatic β-cell function such as the stimulus-secretion coupling and ion channel activity. In addition, we found that, in general, the sensitivity and responses were comparable to those from other in vivo studies reported in the literature. Overall, our results suggest that both systems can serve as effective tools for rapid screening of potential MDC effects on pancreatic β-cell physiology as well for deciphering and better understanding the molecular mechanisms that underlie their action.

## 1. Introduction

Pancreatic β-cells are an endocrine cell type with the unique ability to synthetize, store, and secrete insulin, the only hormone able to decrease circulating blood glucose levels. They are perfectly designed to act as fine-tuned fuel sensors that are stimulated by dietary nutrients, particularly glucose, such that insulin release occurs to ensure appropriate nutrient uptake and storage [1, 2]. The failure of normal pancreatic β-cell function is accepted as not only the hallmark of type 1 and type 2 diabetes, but also an essential component of the pathophysiology in other metabolic diseases like obesity or metabolic syndrome. This β-cell failure may occur in different manners. While in type 1 diabetes, β-cells are destroyed by a β-cell-specific autoimmune process [3], in type 2 diabetes, as well as in obesity-linked type 2 diabetes, β-cells are dysfunctional as they are no longer able to adapt to the elevated insulin demand caused by the systemic insulin resistance [4]. In overweight/obesity conditions, when β-cells are exposed to chronically excess nutrients, insulin secretion initially increases, but eventually β-cell dysfunction and death occur [5]. In addition, the extent of β cell malfunction is known to be correlated with the severity of metabolic syndrome [6].

Genetic predisposition and lifestyle choices are commonly accepted reasons for the occurrence of metabolic disorders [7]. More recently, it has been acknowledged that a certain class of endocrine-disrupting chemicals (EDCs), the so-called metabolism-disrupting chemicals (MDCs), may also promote metabolic disturbances [8–11]. EDCs encompass a heterogeneous group of chemical substances that can interfere with any aspect of the endocrine system, including hormone production, release, transport, metabolism, binding, action, or elimination [8, 12]. The list of EDCs is rapidly growing and includes synthetic chemicals such as plastics, plasticizers, pesticides, industrial solvents, and heavy metals, among others [8]. Despite all the data supporting EDCs role as metabolic disruptors, there is a lack of robust screening methods that allow us to identify potential MDCs leading to disturbances in glucose and lipid metabolism.

This is extremely relevant as the prevalence rates of metabolic diseases have exploded over the last several decades to the extent that nowadays they are considered a global synergy of epidemics. As a matter of fact, about 32.3 million adults were diagnosed with diabetes in the European Union in 2019, up from an estimated 16.8 million adults in 2000, while in America the estimated incidence was 34.2 million [13, 14]. Of no less concern is the prevalence of obesity and overweight. According to the World Health Organization, 39% of adults aged 18 or over were overweight and 13% obese [15]. In turn, the metabolic syndrome is known to increase the risk of type 2 diabetes mellitus fivefold [16].

At present, there are several test guidelines and specific programs for the screening and testing of EDCs. The Organization for Economic Co-operation and Development (OECD) has recently compiled and revised them [17]. Of note, all of them are intended to evaluate estrogenic, androgenic, thyroid hormone, and steroidogenesis pathways. However, other endocrine pathways that may be key to the identification of MDCs are not assayed in the current validated tests. Therefore, there is an urgent need to develop new bioassays to interrogate these endocrine-specific pathways. As pancreatic β-cells are essential for glucose and energy homeostasis, new cell-based assays focused on this specific cell type will contribute to a better identification of potential MDCs as well as to the development of testing strategies for regulatory needs.

The aim of this study was to investigate the effects of a number of model EDCs on pancreatic β- cell survival and function, and to evaluate different pancreatic β-cell models to be used as in vitro test systems. We analyzed seven different chemicals used as: plasticizers, bisphenol-A (BPA), the BPA substitutes, bisphenol-S (BPS) and bisphenol-F (BPF), as well as di(2-ethylhexyl) phthalate (DEHP); pesticides, dichlorodiphenyldichloroethylene (DDE); a representative polyfluoroalkyl compound, perfluorooctanesulfonic acid (PFOS); and the heavy metal cadmium chloride (CdCl_2_). Chemical selection was made based on their widespread use, large production volume, and their potential diabetogenic and obesogenic properties. We explored the effects of these compounds on two different pancreatic β-cell lines: MIN6 (a mouse β-cell line) and EndoC-βH1 (a human β- cell line).

Immortalized rodent β-cell lines have been extensively exploited for the study of β-cell physiology and were demonstrated to be a valuable asset in diabetes research [18]. Nevertheless, they have not been thoroughly evaluated for EDC screening and testing. A limited number of previous studies have looked at the effects of particular EDCs on MIN6 cells; however, to the best of our knowledge, this is the first study assessing the impact of paradigmatic EDCs on a human pancreatic β-cell line. By using these two cell models we compared murine and human β-cell responses, which may be very informative because up-to-date, direct causal relationships between EDCs and metabolic disorders have mainly come from animal studies. Our results highlighted that both β-cell models can offer a consistent and sensitive option for screening putative MDCs.

## 2. Results

### 2.1. Bisphenols A and S alter pancreatic β-cell function in both mouse and human pancreatic β-cells

The effects of BPA on cell viability were first studied in the murine pancreatic β-cell line MIN6 after 24 h of exposure to different concentrations of the chemical ranging from 100 pM to 10 μM. Cell viability was assayed using a combination of three classical dye methods: resazurin (RZ) assay as an indicator of mitochondrial activity, CFDA-AM assay as an indicator of cytoplasmic esterase activity and membrane integrity, and neutral red uptake (NRU) assay, which evaluates lysosomal activity. As illustrated in Figure 1A, cell viability (measured as RZ test) was slightly reduced at the concentration range of 100 pM to 1 μM with the maximum effect at both 10 and 100 nM BPA concentrations (reduction to 87.0 ± 1.4 and 87.9 ± 1.7%, respectively, compared to control (100%)) (Figure 1A). NRU and CFDA-AM assay tests showed a modest decrease in cell viability, which was only significant at 10 nM BPA (reduction to 91.6 ± 1.4 and 88.3 ± 1.7%, respectively, compared to control (100%)) (Figure 1A). The cytotoxic effects of BPA after 48 or 72 h treatment were also explored. In brief, RZ and CFDA-AM assays indicated moderately decreased cell viability at almost all doses tested (1 nM–10 μM) at time point 48 h, while at 72 h after BPA exposure, a more pronounced cytotoxic effect was observed at 10 nM and 10 μM. Lysosomal activity was slightly decreased at 48 h (1 μM BPA) and 72 h (10 nM and 10 μM BPA) (Supplemental Table 1).

**Figure 1.**
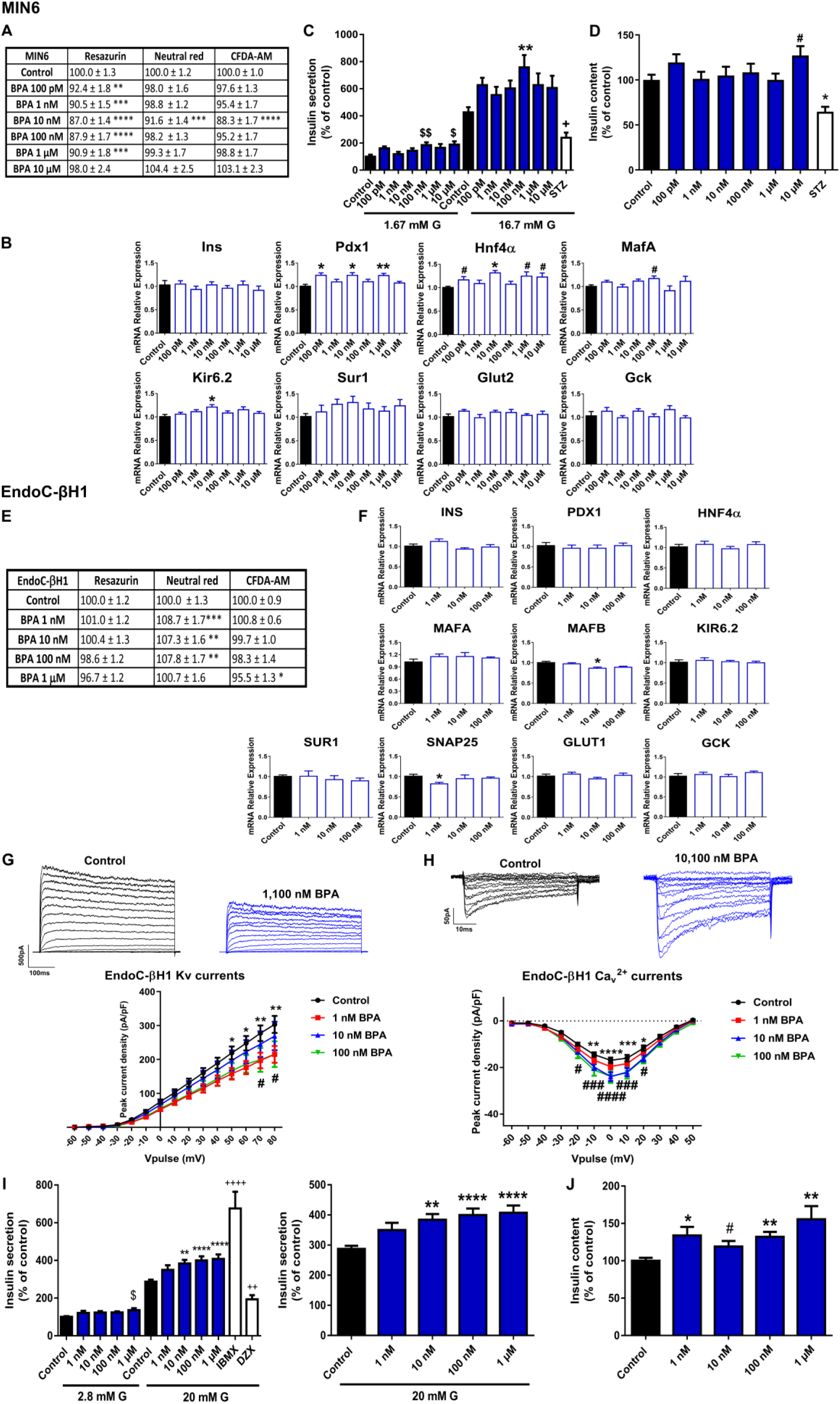
BPA effects on pancreatic β-cells. A) Viability of MIN6 cells treated for 24 h with different BPA concentrations (100 pM–10 μM) as evaluated by RZ, NRU and CFDA-AM assays. n = five independent MafA, Kir6.2, Sur1, Glut2, Gck in MIN6 cells treated for 24 h with different BPA concentrations (100 pM–10 μM). n = three independent experiments. * vs. Control; one-way ANOVA or Kruskal-Wallis. # vs. Control; Student’s t-test. C) Effects of BPA (100 pM–10 μM) on GSIS in MIN6 cells treated for 24 h. n = six independent experiments. * vs. Control 16.7 mM G; two-way ANOVA. $ vs. Control 1.67 mM G and + vs. Control 16.7 mM G; Kruskal-Wallis. D) MIN6 insulin content after 24 h BPA treatment. n = six independent experiments. * vs. Control; Kruskal-Wallis. # vs. Control; Student’s t-test. E) Viability of EndoC-βH1 cells treated for 72 h with different BPA concentrations (1 nM–1 μM) as evaluated by RZ, NRU and CFDA-AM assays. n = four independent experiments. * vs. Control; one-way ANOVA or Kruskal-Wallis. F) mRNA expression of INS, PDX1, HNF4α, MAFA, MAFB, KIR6.2, SUR1, SNAP25, GLUT1 and GCK in EndoC-βH1 cells treated for 72 h with different BPA concentrations (1 nM–100 nM). n = three independent experiments. * vs. Control; Kruskal-Wallis. G) Upper panel, representative recordings of K^+^ currents in response to depolarising voltage pulses in Control or BPA (1 nM–100 nM) EndoC-βH1 treated-cells for 72 h. Lower panel, relationship between K^+^ current density and the voltage of the pulses. Control (n = 12) and BPA (n =12 per condition) cells. * Control vs. 1 nM BPA and # Control vs. 100 nM BPA; two-way ANOVA. H) Upper panel, representative recordings of Ca^2+^ currents in response to depolarising voltage pulses in Control or BPA (1 nM–100 nM) EndoC-βH1 treated-cells for 72 h. Lower panel, relationship between Ca^2+^ current density and the voltage of the pulses. Control (n = 11) and BPA (n =10 per condition) cells. * Control vs. 10 nM BPA and # Control vs. 100 nM BPA ; two-way ANOVA. I) Effects of BPA (1 nM–1 μM) on GSIS in EndoC-βH1 cells treated for 72 h. Left panel: GSIS in response to low glucose (2.8 mM G) and high glucose (20 mM G). Right panel is an inset graph that shows insulin release in response to 20 mM G. n = five independent experiments. * vs. Control 20 mM G; two-way ANOVA. $ vs. Control 2.8 mM G and + vs. Control 20 mM G; one-way ANOVA. J) EndoC-βH1 insulin content after 72 h BPA treatment. n = five independent experiments. * vs. Control; one-way ANOVA. # vs. Control; Student’s t-test. All data are expressed as mean ± SEM. Significance *p < 0.05, **p < 0.01, ***p < 0.001 and ****p < 0.0001; # p < 0.05, ###p < 0.001, and ####p < 0.0001; $p < 0.05, $$p < 0.01; +p < 0.05, ++p < 0.01.

We next analyzed whether BPA treatment for 24 h affected gene expression in MIN6 cells. We focused on the key genes for pancreatic β-cell function and identity. We found that Pdx1 and Hnf4α gene expressions were upregulated at various BPA concentrations (100 pM, 10 nM, and 1 μM) (Figure 1B); Hnf4α was elevated at 10 μM as well. BPA treatment also promoted increased expression of MafA and Kir6.2, at 100 and 10 nM concentrations, respectively (Figure 1B). These results were consistent with the increments of glucose-stimulated insulin secretion (GSIS) found in the BPA-treated cells compared to controls (Figure 1C). The increase was only statistically significant at 100 nM BPA, although a similar trend was observed at all tested doses (Figure 1C). In addition, insulin secretion in response to low glucose was also increased at 100 nM and 10 μM BPA. The insulin content was found to be increased at the highest dose tested (10 μM) (Figure 1D).

EndoC-βH1 cell viability was not reduced after 24 h (Supplemental Table 2), 48 h (Supplemental Table 2), or 72 h (Figure 1E) BPA exposure. Unlike the results found in the murine cell model, only a slight decrease in mitochondrial activity (1 μM BPA) and membrane integrity (100 nM and 1 μM BPA) at 48 h was observed (Supplemental Table 2). On the contrary, BPA treatment for 72 h (1, 10, and 100 nM) modestly augmented the percentage of lysosomal activity (NRU assay) (Figure 1E). No effect was observed on GSIS when cells were treated for 24 or 48 h with BPA (Supplemental Figures 1A and 1B, respectively); however, a marked dose-dependent increase in GSIS was observed in EndoC-βH1 cells treated with BPA for 72 h compared to controls (Figure 1I). This effect was statistically significant at 10, 100 nM and 1 μM doses. Basal insulin secretion was also increased at 1 μM BPA. In a similar manner, increased insulin content was found in the BPA-treated cells (72 h), which was significant at all doses tested (Figure 1J). The impact of BPA on insulin release in EndoC-βH1 cells was not related to changes in gene expression (Figure 1F) but to a disruptive effect on the electrical activity (Figures 1G and 1H). Voltage-gated K^+^ and Ca^2+^ currents were recorded in human pancreatic β-cells treated with vehicle or BPA for 72 h. As shown in Figure 1G, BPA exposure resulted in decreased K^+^ currents at both 1 and 100 nM concentrations. In addition, augmented Ca^2+^ currents were found in BPA-treated cells (10 and 100 nM) (Figure 1H).

BPS was also studied since this compound has become a common BPA substitute. Like BPA, BPS promoted a modest decrease in cell viability (measured as RZ test) in the pancreatic β-cell line MIN6 after 24 h treatment at all doses tested (Figure 2A). This effect was also observed at 48 h, although it was less pronounced at 72 h (Supplemental Table 1). The CFDA-AM assay also showed a slight but significant cytotoxic effect of BPS in MIN6 cells treated for 24 (1 nM–10 μM) (Figure 2A), 48, or 72 h (10 nM–10 μM) (Supplemental Table 1). The NRU assay manifested a much more limited BPS effect, which was only significant at 48 h (100 pM and 10 μM) (Supplemental Table 1). These data indicated that BPS slightly impaired mitochondrial activity and membrane integrity, but it did not affect lysosomal activity.

**Figure 2.**
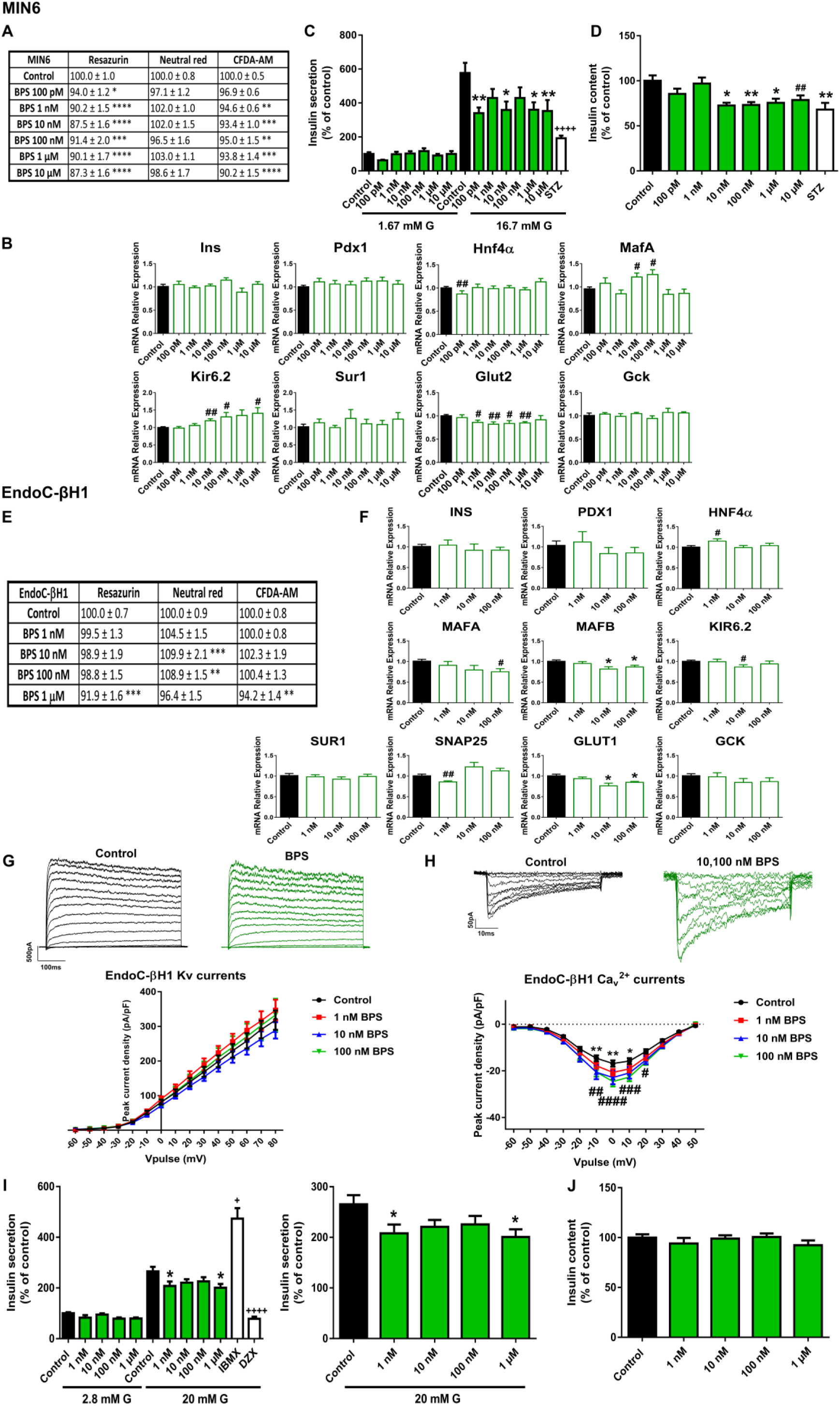
BPS effects on pancreatic β-cells. A) Viability of MIN6 cells treated for 24 h with different BPS MafA, Kir6.2, Sur1, Glut2, Gck in MIN6 cells treated for 24 h with different BPS concentrations (100 pM–10 μM). n = three independent experiments. # vs. Control; Student’s t-test. C) Effects of BPS (100 pM–10 μM) on GSIS in MIN6 cells treated for 24 h. n = five independent experiments. * vs. Control 16.7 mM G; two-way ANOVA. + vs. Control 16.7 mM G; Kruskal-Wallis. D) MIN6 insulin content after 24 h BPS treatment. n = five independent experiments. * vs. Control; one-way ANOVA. # vs. Control; Student’s t-test. E) Viability of EndoC-βH1 cells treated for 48 h with different BPS concentrations (1 nM–1 μM) as evaluated by RZ, NRU and CFDA-AM assays. n = four independent experiments. * vs. Control; one-way ANOVA. F) mRNA expression of INS, PDX1, HNF4α, MAFA, MAFB, KIR6.2, SUR1, SNAP25, GLUT1 and GCK in EndoC-βH1 cells treated for 48 h with different BPS concentrations (1 nM–100 nM). n = three independent experiments.* vs. Control; Kruskal-Wallis. # vs. Control; Student’s t-test. G) Upper panel, representative recordings of K^+^ currents in response to depolarising voltage pulses in Control or BPS (1 nM–100 nM) EndoC-βH1 treated- cells for 48 h. Lower panel, relationship between K^+^ current density and the voltage of the pulses. Control (n = 27) and BPS (n = 13–20 per condition) cells. H) Upper panel, representative recordings of Ca^2+^ currents in response to depolarising voltage pulses in Control or BPS (1 nM–100 nM) EndoC-βH1 treated-cells for 48 h. Lower panel, relationship between Ca^2+^ current density and the voltage of the pulses. Control (n = 12) and BPS (n =11–12 per condition) cells. * Control vs. 10 nM BPS and # Control vs. 100 nM BPS; two-way ANOVA. I) Effects of BPS (1 nM–1 μM) on GSIS in EndoC-βH1 cells treated for 48 h. Left panel: GSIS in response to low glucose (2.8 mM G) and high glucose (20 mM G). Right panel is an inset graph that shows insulin release in response to 20 mM G. n = five independent experiments. * vs. Control 20 mM G; two-way ANOVA. + vs. Control 20 mM G; Kruskal-Wallis. J) EndoC-βH1 insulin content after 48 h BPS treatment. n = five independent experiments. All data are expressed as mean ± SEM. Significance *p < 0.05, **p < 0.01, ***p < 0.001 and ****p < 0.0001; # p < 0.05, ## p < 0.01, ###p < 0.001, and ####p < 0.0001; ++++p < 0.0001.

Changes in the gene expression profile were also explored in the murine cells treated with BPS for 24 h (Figure 2B). The most remarkable effect was a marked decrease in Glut2 gene expression in a wide range of doses from 1 nM to 1 μM. Decreased expression of Hnf4α was also found, although it was only statistically significant at the lowest dose (100 pM). By contrast, an upregulation of MafA (10 and 100 nM) and Kir6.2 (10, 100 nM, and 10 μM) was quantified. Together with a reduction effect on the glucose transporter expression we found markedly diminished GSIS at all doses tested, which was statistically different from controls at 100 pM, 10 nM, 1 and 10 μM BPS concentrations (Figure 2C). Parallel to this effect on insulin secretion, we found that BPS-treated cells exhibited a reduction in insulin content at 10, 100 nM, 1 and 10 μM concentrations (Figure 2D).

Then, we analyzed BPS effects on the human cellular model EndoC-βH1. As in the case of BPA, BPS did not affect viability in EndoC-βH1 cells. However, a modest increase in the lysosomal activity (NRU) after 24 h (10 nM) (Supplemental Table 2), 48 h (10 and 100 nM) (Figure 2E), and 72 h (100 nM) treatments was found (Supplemental Table 2).

Similar to the effects reported in MIN6, 48 h BPS treatment resulted in a marked decreased expression of the gene encoding for the main glucose transporter (GLUT1) in EndoC-βH1 cells. The effect was found to be statistically significant at 10 and 100 nM concentrations (Figure 2F). In addition, decreased expressions of MAFA (100 nM), MAFB (10 and 100 nM), SNAP25 (1 nM), and KIR6.2 (10 nM) were quantified (Figure 2F). These changes were correlated with a marked diminished insulin release at the same time point of 48 h. A clear decreasing trend was observed at all doses tested although the statistically significant effects were found at 1 nM and 1 μM BPS doses (Figure 2I). No effects on insulin content were reported (Figure 2J). As regards electrical activity, no effects on K^+^ currents were quantified (Figure 2G), although increased calcium currents were registered in response to 10 and 100 nM BPS (Figure 2H).

The second most common BPA substitute, BPF, was also included in the study. For BPF we did not find any effect on GSIS in EndoC-βH1-treated cells compared to controls (Supplemental Figure 2A) at any of the doses tested. A slight decrease in cell viability (measured as RZ assay) was found with the most significant effects after 24 and 48 h treatments. Membrane integrity was also slightly reduced after 24 h treatment, while lysosomal activity showed a slight increase (Supplemental Figure 2B).

### 2.2 The phthalate DEHP disrupts murine and human pancreatic β-cell function in a similar manner

As shown in Figure 3A, RZ and NRU tests showed no effect of DEHP on cell viability in MIN6 cells treated for 24 h at most doses tested, while the CFDA-AM assay manifested a modest decrease in membrane integrity in response to DEHP (10 nM–10 μM). Of note, after 48 h treatment there was a consistent decrease in cell viability at the highest dose of DEHP (10 μM) as measured in both NRU (79.5 ± 1.7%) and CFDA-AM (83.2 ± 3.2%) assays. The effect was even more pronounced at 72 h, when a dramatic decrease in the percentage of cell viability was quantified in all the assays performed: RZ (59.1 ± 3.7%), NRU (47.1 ± 2.9%) and CFDA-AM (65.9 ± 3.3%) (Supplemental Table 1).

**Figure 3.**
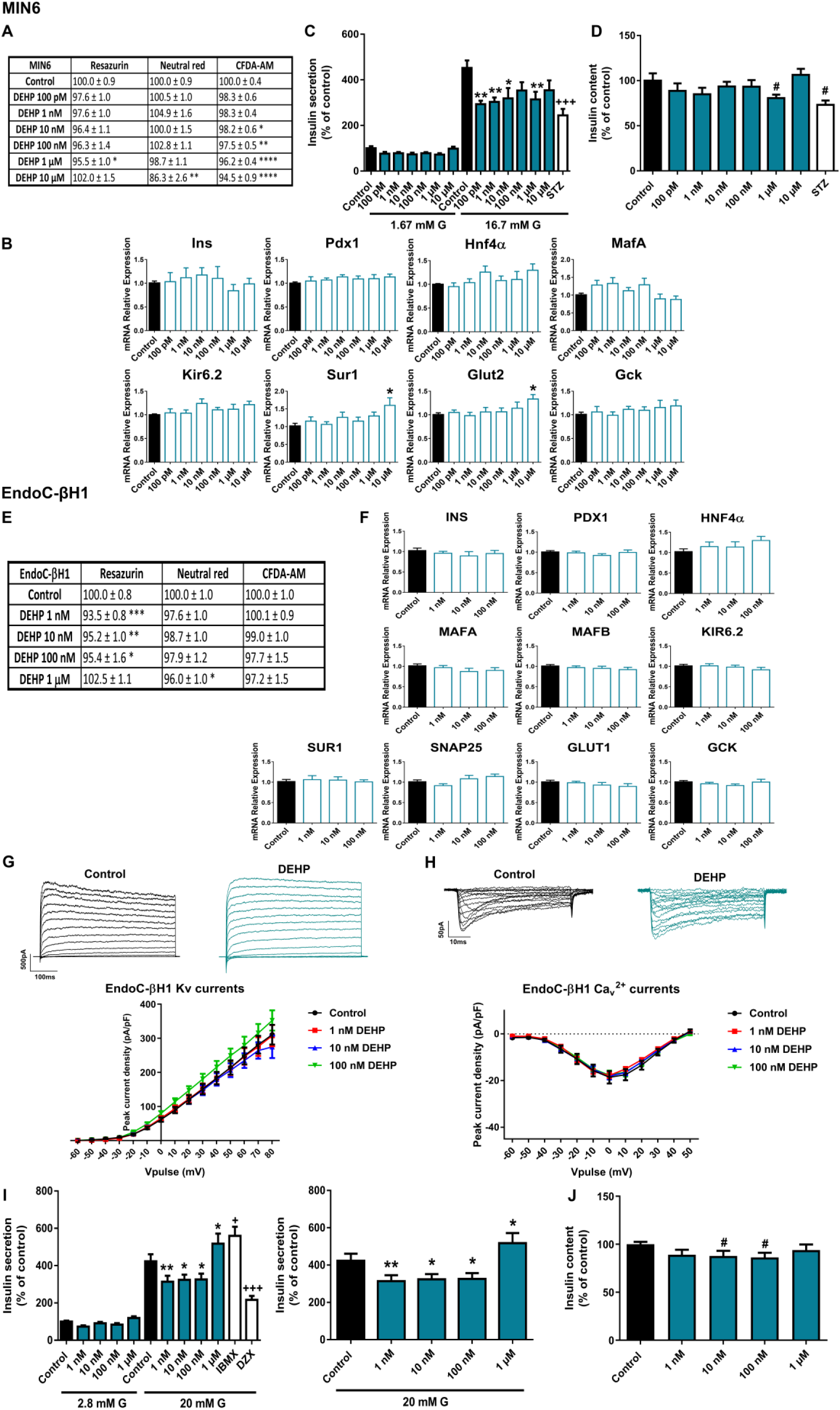
DEHP effects on pancreatic β-cells. A) Viability of MIN6 cells treated for 24 h with different DEHP experiments. * vs. Control; Kruskal-Wallis. B) mRNA expression of Ins, Pdx1, Hnf4α, MafA, Kir6.2, Sur1, Glut2, Gck in MIN6 cells treated for 24 h with different DEHP concentrations (100 pM–10 μM). n = five independent experiments. * vs. Control; Kruskal-Wallis. C) Effects of DEHP (100 pM–10 μM) on GSIS in MIN6 cells treated for 24 h. n = four independent experiments. * vs. Control 16.7 mM G; two-way ANOVA. + vs. Control 16.7 mM G; one-way ANOVA. D) MIN6 insulin content after 24 h DEHP treatment. n = four independent experiments. # vs. Control; Student’s t-test. E) Viability of EndoC-βH1 cells treated for 7 d with different DEHP concentrations (1 nM–1 μM) as evaluated by RZ, NRU and CFDA-AM assays. n = four independent experiments. * vs. Control; one-way ANOVA. F) mRNA expression of INS, PDX1, HNF4α, MAFA, MAFB, KIR6.2, SUR1, SNAP25, GLUT1 and GCK in EndoC-βH1 cells treated for 7 d with different DEHP concentrations (1 nM–100 nM). n = four independent experiments. G) Upper panel, representative recordings of K^+^ currents in response to depolarising voltage pulses in Control or DEHP (1 nM–100 nM) EndoC-βH1 treated-cells for 7 d. Lower panel, relationship between K^+^ current density and the voltage of the pulses. Control (n = 15) and DEHP (n = 12–15 per condition) cells. H) Upper panel, representative recordings of Ca^2+^ currents in response to depolarising voltage pulses in Control or DEHP (1 nM–100 nM) EndoC-βH1 treated-cells for 7 d. Lower panel, relationship between Ca^2+^ current density and the voltage of the pulses. Control (n = 11) and DEHP (n = 11 per condition) cells. I) Effects of DEHP (1 nM–1 μM) on GSIS in EndoC-βH1 cells treated for 7 d. Left panel: GSIS in response to low glucose (2.8 mM G) and high glucose (20 mM G). Right panel is an inset graph that shows insulin release in response to 20 mM G. n = five independent experiments. * vs. Control 20 mM G; two-way ANOVA. + vs. Control 20 mM G; Kruskal-Wallis. J) EndoC-βH1 insulin content after 7 d DEHP treatment. n = five independent experiments. # vs. Control; Student’s t-test. All data are expressed as mean ± SEM. Significance *p < 0.05, **p < 0.01, ***p < 0.001 and ****p < 0.0001; # p < 0.05 ; +p < 0.05, +++p < 0.001.

No significant changes were found at the level of gene expression in response to 24 h DEHP treatment except for an increase in the expression of Sur1 and Glut2 genes at the highest dose tested (10 μM) (Figure 3B). However, at the functional level, we observed an impairment of insulin release in response to stimulatory glucose concentration in DEHP-treated cells for 24 h compared to controls (Figure 3C). The effect was statistically significant at almost all doses tested. Decreased insulin content was also observed at the highest dose (1 μM) (Figure 3D). Although the disruptive effect of DEHP on secretion was similar in human and murine cells, we observed a clear difference in the exposure time required to achieve this effect. We did not find any change in insulin secretion in EndoC-βH1 after 72 h DEHP treatment (Supplemental Figure 3), so we extended our study to 7 d. As shown in Figure 3I, we found that 7 d treatment with DEHP resulted in a marked decrease in GSIS at 1, 10, and 100 nM concentrations. Curiously, at the highest dose tested, 1 μM, DEHP-treated cells manifested increased insulin release (Figure 3I). In addition to that, a decrease in insulin content in response to 10 and 100 nM DEHP exposure was quantified (Figure 3J). No changes at the level of gene expression (Figure 3F), K^+^ or Ca^2^ currents (Figures 3G and 3H) were observed at this time point. The effects of DEHP on cell viability were explored after 24, 48, 72 h (Supplemental Table 2) and 7 d of exposure (Figure 3E). A modest decrease in membrane integrity was observed at the 24, 48, and 72 h time points in response to the highest DEHP dose tested (1 μM). In addition, after 7 d treatment, mitochondrial activity was decreased at the tested concentration range of 1 to 100 nM, while lysosomal activity was found to be decreased only at the highest dose (1 μM) (Figure 3E).

### 2.3. Different effects of PFOS exposure on human and murine pancreatic β-cell models

Slight inhibition of MIN6 cell viability was detected after 24 h PFOS exposure (Figure 4A). RZ reduction and membrane integrity (CFDA-AM assay) were the most sensitive cellular processes affected, with a reduction to approximately 87–92% compared to controls (100%) in response to all tested doses. Lysosomal activity was less affected (reduction to 90–96%), with no effect at 100 pM and 1 μM doses (Figure 4A). The 3 metabolic assays showed similar results of reduced viability after 48 h treatment with more significant differences at 10 nM and 1 μM. On the contrary, no significant changes in viability were observed after 72 h treatment (Supplemental Table 1). PFOS exposure also induced a reduction in Glut2 gene expression at almost all the doses analyzed (10 nM–10 μM) as well as a sharp reduction in MafA gene expression at 1 μM dose (Figure 4B). In line with this, insulin release in response to stimulatory glucose concentration was markedly reduced in 24 h PFOS-treated cells at 100 pM, 100 nM, and 10 μM concentrations (Figure 4C). No effects were found on insulin content (Figure 4D).

**Figure 4.**
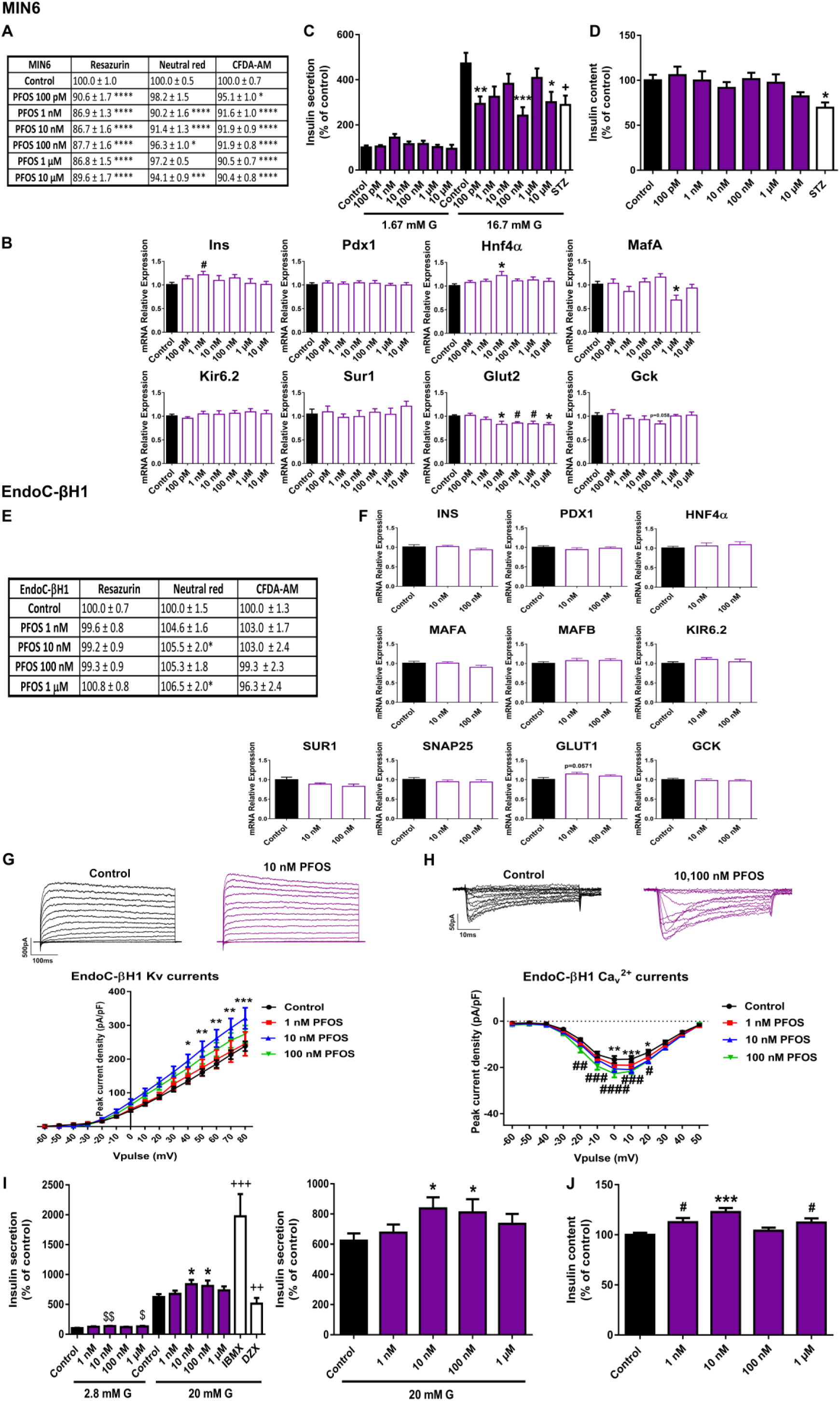
PFOS effects on pancreatic β-cells. A) Viability of MIN6 cells treated for 24 h with different PFOS MafA, Kir6.2, Sur1, Glut2, Gck in MIN6 cells treated for 24 h with different PFOS concentrations (100 pM– 10 μM). n = three independent experiments. * vs. Control; one-way ANOVA. #vs. Control; Student’s t-test. C) Effects of PFOS (100 pM–10 μM) on GSIS in MIN6 cells treated for 24 h. n =five independent experiments. vs. Control 16.7 mM G; two-way ANOVA. + vs. Control 16.7 mM G; one-way ANOVA. D) MIN6 insulin content after 24 h PFOS treatment. n = five independent experiments. * vs. Control; Kruskal-Wallis. E) Viability of EndoC-βH1 cells treated for 72 h with different PFOS concentrations (1 nM–1 μM) as evaluated by RZ, NRU and CFDA-AM assays. n = five independent experiments. * vs. Control; Kruskal-Wallis. F) mRNA expression of INS, PDX1, HNF4α, MAFA, MAFB, KIR6.2, SUR1, SNAP25, GLUT1 and GCK in EndoC-βH1 cells treated for 72 h with different PFOS concentrations (1 nM–100 nM). n = three independent experiments. G) Upper panel, representative recordings of K^+^ currents in response to depolarising voltage pulses in Control or PFOS (1 nM–100 nM) EndoC-βH1 treated-cells for 72 h. Lower panel, relationship between K^+^ current density and the voltage of the pulses. Control (n = 11) and PFOS (n = 11–14 per condition) cells. * Control vs. 10 nM PFOS; two-way ANOVA. H) Upper panel, representative recordings of Ca^2+^ currents in response to depolarising voltage pulses in Control or PFOS (1 nM–100 nM) EndoC-βH1 treated- cells for 72 h. Lower panel, relationship between Ca^2+^ current density and the voltage of the pulses. Control (n = 10) and PFOS (n = 10–12 per condition) cells. * Control vs. 10 nM PFOS and # Control vs. 100 nM PFOS; two-way ANOVA. I) Effects of PFOS (1 nM–1 μM) on GSIS in EndoC-βH1 cells treated for 72 h. Left panel: GSIS in response to low glucose (2.8 mM G) and high glucose (20 mM G). Right panel is an inset graph that shows insulin release in response to 20 mM G. n = five independent experiments. * vs. Control 20 mM G; two-way ANOVA. $ vs. Control 2.8 mM G and + vs. Control 20 mM G; Kruskal-Wallis. J) EndoC-βH1 insulin content after 72 h PFOS treatment. n = five independent experiments. * vs. Control; Kruskal-Wallis. # vs. Control; Student’s t-test. All data are expressed as mean ± SEM. Significance *p < 0.05, **p < 0.01, ***p < 0.001 and ****p < 0.0001; # p < 0.05, ##p < 0.01, ###p < 0.001, and ####p < 0.0001; $p < 0.05, $$p < 0.01; +p < 0.05, ++p < 0.01, +++p < 0.001.

In human pancreatic β-cells we found quite different results. No remarkable changes in cell viability were observed at any of the time points assayed; instead, a slight increase in lysosomal activity (NRU assay) after 24 and 48 h (1 nM–1 μM) (Supplemental Table 2) and 72 h treatment (10 nM and 1 μM) was reported (Figure 4E). No significant changes were observed at the level of gene expression (Figure 4F). However, a consistent tendency toward increased expression of the glucose transporter GLUT1 was observed (p=0.0571) (Figure 4F). As shown in Figure 4I, an increase in GSIS was quantified at 10 and 100 nM PFOS doses as well as in basal insulin secretion (10 nM and 1 μM). In line with this, increased insulin content at 1, 10 nM, and 1 μM PFOS was found (Figure 4J). Importantly, this increase in the secretory capacity of the human β-cells was accompanied by an increase in K^+^ currents at 10 nM (Figure 4G) and Ca^2+^ currents at 10 and 100 nM PFOS (Figure 4H).

### 2.4. CdCl_2_ effects on pancreatic β-cell function and viability

CdCl_2_ was the EDC that caused the greatest inhibition of MIN6 cell viability. As shown in Figure 5A, 24 h CdCl_2_ treatment significantly affected RZ reduction (100 pM–10 μM), NRU (100 nM–10 μM) and CFDA-AM (1 nM–10 μM). Similar results were found after 48 and 72 h treatment (Supplemental Table 1). The effects were relatively modest at concentrations in the range of 100 pM to 1 μM; however, all three metabolic tests assayed showed a very pronounced cytotoxic effect of CdCl_2_ at the highest dose (10 μM) (RZ 22.0 ± 2.1%, NRU 12.2 ± 2.2%, CFDA-AM 39.9 ± 1.1%), which was maintained at all-time points analyzed. This dose was excluded from the study.

**Figure 5.**
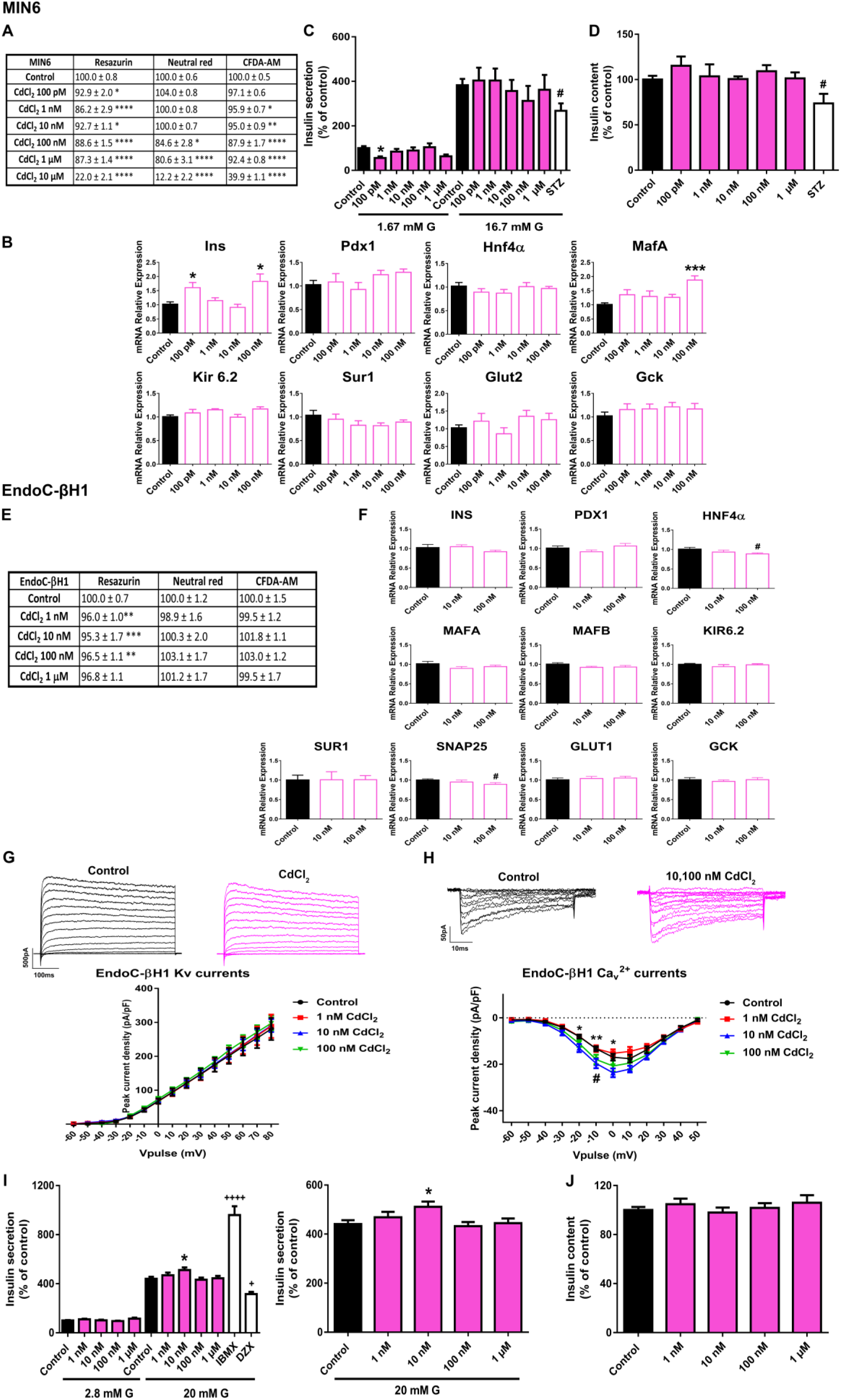
CdCl2 effects on pancreatic β-cells. A) Viability of MIN6 cells treated for 24 h with different CdCl2 MafA, Kir6.2, Sur1, Glut2, Gck in MIN6 cells treated for 24 h with different CdCl2 concentrations (100 pM– 100 nM). n = three independent experiments. * vs. Control; one-way ANOVA. C) Effects of CdCl2 (100 pM– 1 μM) on GSIS in MIN6 cells treated for 24 h. n = three independent experiments. * vs. Control 1.67 mM G; two-way ANOVA. # vs. Control 16.7 mM G; Student’s t-test. D) MIN6 insulin content after 24 h CdCl2 treatment. n = three independent experiments. # vs. Control; Student’s t-test. E) Viability of EndoC-βH1 cells treated for 72 h with different CdCl2 concentrations (1 nM–1 μM) as evaluated by RZ, NRU and CFDA-AM assays. n = five independent experiments. * vs. Control; Kruskal-Wallis. F) mRNA expression of INS, PDX1, HNF4α, MAFA, MAFB, KIR6.2, SUR1, SNAP25, GLUT1 and GCK in EndoC-βH1 cells treated for 72 h with different CdCl2 concentrations (10, 100 nM). n = three independent experiments. # vs. Control; Student’s t- test. G) Upper panel, representative recordings of K^+^ currents in response to depolarising voltage pulses in Control or CdCl2 (1 nM–100 nM) EndoC-βH1 treated-cells for 72 h. Lower panel, relationship between K^+^ current density and the voltage of the pulses. Control (n = 10) and CdCl2 (n = 10 per condition) cells. H) Upper panel, representative recordings of Ca^2+^ currents in response to depolarising voltage pulses in Control or CdCl2 (1 nM–100 nM) EndoC-βH1 treated-cells for 72 h. Lower panel, relationship between Ca^2+^ current density and the voltage of the pulses. Control (n = 8) and CdCl2 (n = 8–10 per condition) cells. * Control vs. 10 nM CdCl2 and # Control vs. 100 nM CdCl2; two-way ANOVA. I) Effects of CdCl2 (1 nM–1 μM) on GSIS in EndoC-βH1 cells treated for 72 h. Left panel: GSIS in response to low glucose (2.8 mM G) and high glucose (20 mM G). Right panel is an inset graph that shows insulin release in response to 20 mM G . n = five independent experiments. * vs. Control 20 mM G; two-way ANOVA. + vs. Control 20 mM G; one-way ANOVA. J) EndoC-βH1 insulin content after 72 h CdCl2 treatment. n = five independent experiments. All data are expressed as mean ± SEM. Significance *p < 0.05, **p < 0.01, ***p < 0.001 and ****p < 0.0001; #p < 0.05; +p < 0.05, ++++p < 0.0001.

When cells were treated with CdCl_2_ for 24 h, significant increased expressions of Ins (100 pM and 100 nM) and MafA (100 nM) genes were quantified (Figure 5B). Nevertheless, this was not related to any effect on GSIS (Figure 5C) or insulin content (Figure 5D).

In EndoC-βH1 cells, the cytotoxic effect of CdCl_2_ was clearly less pronounced. Only a slight decrease in cell viability (measured as RZ reduction) at 48 h (1nM–1 μM) (Supplemental Table 2) and 72 h (1–100 nM) (Figure 5E) was found. A modest decrease in HNF4α and SNAP25 gene expressions at 100 nM dose was found (Figure 5F). Regarding changes in electrical activity, enhanced Ca^2+^ currents at 10 and 100 nM CdCl_2_ doses compared to control were noted (Figure 5H), but no effects on K^+^ currents (Fig. 5G) were found. In addition, augmented GSIS was observed in CdCl_2_-treated cells at 10 nM (Figure 5I). No effects on insulin content were reported (Figure 5J).

### 2.5. DDE exposure did not compromise pancreatic β-cell function or viability

MIN6 cells treated with DDE for 24 h exhibited a modest decrease in viability measured as RZ reduction (10 nM) and membrane integrity (10 nM, 1 and 10 μM) (Figure 6A). However, this effect was not confirmed in the NRU assay (Figure 6A). A more pronounced reduction was found after 48 h of DDE exposure in the three assays performed at 10 nM (RZ 85.4 ± 1.8%, NRU 89.1 ± 1.9%, CFDA-AM 93.5 ± 1.6%) and 10 μM (RZ 77.4 ± 4.1%, NRU 79.5 ± 4.5%, CFDA-AM 87.6 ± 2.2%) concentrations (Supplemental Table 1). At 72 h, a clear cytotoxic effect of DDE was found at the highest dose assayed (10 μM) (RZ 40.7 ± 2.4%, CFDA-AM 69.9 ± 5.1%) (Supplemental Table 1).

**Figure 6.**
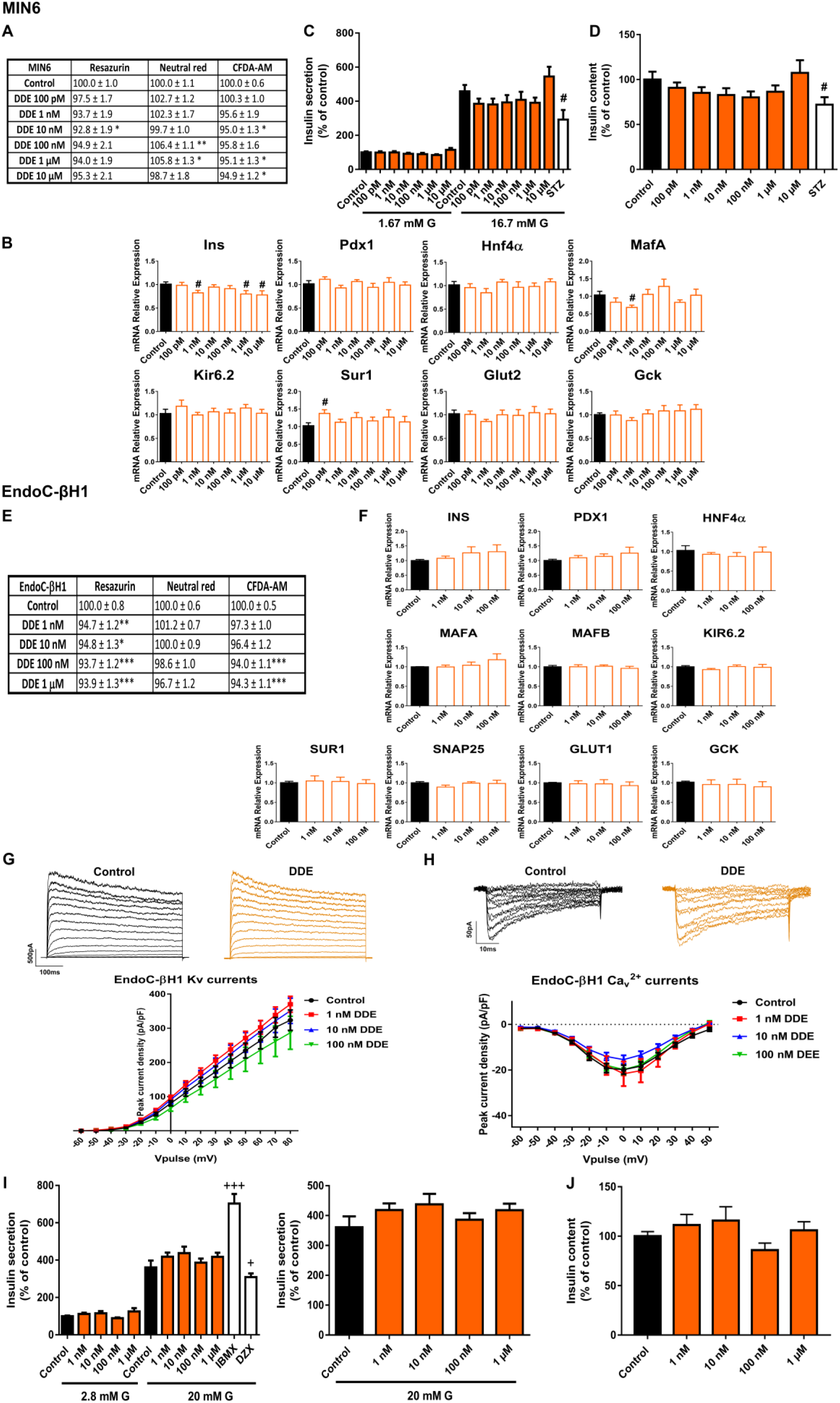
DDE effects on pancreatic β-cells. A) Viability of MIN6 cells treated for 24 h with different DDE concentrations (100 pM–10 μM) as evaluated by RZ, NRU and CFDA-AM assays. n = six-seven independentMafA, Kir6.2, Sur1, Glut2, Gck in MIN6 cells treated for 24 h with different DDE concentrations (100 pM–10 μM). n = three independent experiments. # vs. Control; Student’s t-test. C) Effects of DDE (100 pM–10 μM) on GSIS in MIN6 cells treated for 24 h. n = four independent experiments. # vs. Control; Student’s t-test. D) MIN6 insulin content after 24 h DDE treatment. n= three independent experiments. # vs. Control; Student’s t-test. E) Viability of EndoC-βH1 cells treated for 7 d with different DDE concentrations (1 nM–1 μM) as evaluated by RZ, NRU and CFDA-AM assays. n= eight independent experiments. * vs. Control; one-way ANOVA or Kruskal-Wallis . F) mRNA expression of INS, PDX1, HNF4α, MAFA, MAFB, KIR6.2, SUR1, SNAP25, GLUT1 and GCK in EndoC-βH1 cells treated for 7 d with different DDE concentrations (1 nM–100 nM). n = four independent experiments. G) Upper panel, representative recordings of K^+^ currents in response to depolarising voltage pulses in Control or DDE (1 nM–100 nM) EndoC-βH1 treated-cells for 7 d. Lower panel, relationship between K^+^ current density and the voltage of the pulses. Control (n = 9) and DDE (n = 10 per condition) cells. H) Upper panel, representative recordings of Ca^2+^ currents in response to depolarising voltage pulses in Control or DDE (1 nM–100 nM) EndoC-βH1 treated-cells for 7 d. Lower panel, relationship between Ca^2+^ current density and the voltage of the pulses. Control (n = 8) and DDE (n = 9–10 per condition) cells. I) Effects of DDE (1 nM−1 μM) on GSIS in EndoC-βH1 cells treated for 7 d. Left panel: GSIS in response to low glucose (2.8 mM G) and high glucose (20 mM G). Right panel is an inset graph that shows insulin release in response to 20 mM G. n = three independent experiments. +vs. Control 20 mM G; Kruskal-Wallis. J) EndoC-βH1 insulin content after 7 d DDE treatment. n = three independent experiments. All data are expressed as mean ± SEM. Significance *p < 0.05, **p < 0.01, ***p < 0.001; # p < 0.05; +p < 0.05, +++p < 0.001.

At the level of gene expression, the most profound change detected was a reduction in Ins gene expression at 1 nM, 1 and 10 μM DDE doses (24 h treatment). Furthermore, diminished expression of the MafA gene was found at 1 nM DDE (Figure 6B). GSIS and content was quantified after 24 h treatment with DDE. Both parameters showed a tendency to decrease at DDE doses in the range from 100 pM to 1 μM; however, this finding did not reach statistical significance (Figures 6C and 6D).

Insulin secretion capacity of EndoC-βH1 was not affected by DDE treatment at short periods of 72 h (Supplemental Figure 4) or at longer times of 7 d (Figure 6I). No changes in insulin content were observed either (Figure 6J). A modest decrease in mitochondrial activity and membrane integrity after 24 h (100 nM and 1 μM) and 48 h DDE treatment (1 μM) was quantified (Supplemental Table 2), while no significant changes in cell viability were observed at 72 h (Supplemental Table 2). 7 d of DDE exposure caused a very slight decrease in mitochondrial activity (reduction to approximately 95%, compared to control (100%)) at all doses tested in the nanomolar range (Figure 6E). Finally, electrophysiological recordings of K^+^ and Ca^2+^ currents in DDE-treated β-cells did not show any differences compared to control condition at any of the doses tested (Figures 6G and 6H).

## 3. Discussion

In the present study, we evaluated the effects of selected EDCs within a range of doses relevant to human exposure on important parameters for pancreatic β-cell function. It is important to note that, with the exception of high doses of CdCl_2_, the concentrations tested did not display significant cytotoxic effects either in murine or in human β-cells. In all cases, the viability was above 80%, which highlights that the adverse effects of EDCs reported here could not be attributed to factors such as cytotoxicity.

Voltage-gated ion channels in pancreatic β-cells, such as K^+^ and Ca^2+^ channels, are a key part of the molecular pathway triggering insulin secretion. This is because the stimulation of insulin release by glucose is linked to membrane depolarization and action potential generation. In short, glucose enters β-cells through the glucose transporter Glut2 in mice, and GLUT1 in humans.

Then, glucose is metabolized, leading to an elevation of the cytoplasmic ATP/ADP ratio, which promotes the closure of the ATP-sensitive K^+^ channels (K_ATP_) and K^+^ efflux reduction. This causes membrane depolarization, opening of voltage-dependent Ca^2+^ channels and TTX-sensitive Na^+^ channels, and elevation of cytosolic free Ca^2+^ concentration, which ultimately stimulates the exocytosis of insulin-containing granules [19]. Finally, delayed rectifying voltage-dependent potassium channels (Kv) and calcium-sensitive voltage-dependent K^+^ channels repolarize the membrane, reducing insulin release [20]. Thus, any alteration affecting ion channel activity may lead to altered pancreatic β-cell function.

Here, we evaluated for the first time whether some paradigmatic EDCs may impair glucose-induced electrical activity in a functional human β-cell line. This is highly relevant as human-translatable models have been revealed as essential for a complete understanding of the mechanisms underlying pancreatic β-cell failure in diabetes and other metabolic disorders. In keeping with this, the human β-cell line EndoC-βH1 was recently established as a robust and valid model for studying human β-cell physiology and also for the screening of potential drug target candidates [21, 22]. Furthermore, it was demonstrated that this cellular model expresses ion channel counterparts with characteristics similar to those found in human β-cells, acknowledging that this is a valuable model comparable with primary human β-cells [23].

We found that some of the selected EDCs impaired global K^+^ and/or Ca^2+^ currents. In particular, we observed that BPA treatment led to reduced K^+^ currents in EndoC-βH1 cells, a finding that agreed with previously described results in mouse β-cells, and which was considered as dependent on the activation of the estrogen receptor β [24]. Voltage-gated K^+^ currents in human pancreatic β-cells include two components. The first one is driven by large-conductance Ca^2+^- activated K^+^ channels (big K^+^ channels or BK channels), as it is activated in a rapid manner upon membrane depolarization, depends on Ca^2+^ influx, and is inhibited by iberiotoxin. The second one relies on delayed rectifying K^+^ channels (Kv2.2 and Kv1.6) [25]. Whether BPA directly affects the specific activity of BK channels, Kv channels, or both, in human β-cells requires further investigation. In any case, the inhibition of K^+^ channel activity is known to promote increased insulin secretion [25, 26], an effect that is mimicked by BPA.

Another major finding here reported is that low doses of BPA, PFOS, and CdCl_2_ augmented Ca^2+^ currents in human β-cells. As insulin granule exocytosis is evoked by the rise in [Ca^2+^]_i_ [27], it is tempting to speculate that the modulation of voltage-gated Ca^2+^ channels elicited by the abovementioned EDCs may be responsible, at least in part, for the upregulation of insulin secretion found. Of note, BPS treatment also selectively enhanced Ca^2+^currents, although this effect did not correlate with an increase but rather a decrease in insulin secretion. One possible explanation is that Ca^2+^ channel activity upregulation might be an associated compensatory response to the reduced β-cell insulin release as it happens in aged pancreatic β-cells from mice with lower insulin sensitivity [28]. It may also be the case that BPS is negatively regulating the exocytosis machinery.

Clinical relevance of adequate insulin release in response to circulating blood glucose levels is unquestionable since insulin is the only hormone able to decrease glycemia levels. Equally important is that this occurs in the appropriate range since both too low and too high insulin levels are harmful for the physiological function of body systems. On one hand, sustained impaired insulin secretion will cause hyperglycemia and, eventually, diabetes. On the other hand, an excess of circulating insulin levels will promote hyperinsulinemia and, as a consequence, systemic insulin resistance, a key feature of type 2 diabetes [29].

For clarity of discussion, results of insulin secretion and content for the different EDC categories will be separately dissected. In addition, a comparison between murine and human in vitro models as well as correlation with the in vivo studies in the literature will be addressed.

### Bisphenols

We found that BPA in the dose range of 1 nM to 1 μM markedly increased insulin secretion in both human (EndoC-βH1) and murine (MIN6) pancreatic β-cell lines. This finding agreed with previous published results in mouse [30–32] and human isolated islets [33] and also in other murine pancreatic β-cell lines [34]. It was also known that BPA treatment led to augmented insulin content in mouse islet β-cells [31], an effect that was reproduced here in the human β-cell EndoC-βH1. These results correlated well with the in vivo response elicited by BPA exposure as BPA-treated animals manifested high plasma insulin concentration and insulin resistance, both hallmarks of prediabetes [30]. Epidemiological studies in humans have largely pointed in the same direction [8, 35, 36]. Our data suggested the important role of Ca^2+^ and K^+^ channel activity rather than genetic modification as the molecular mechanism underlying BPA action in pancreatic β-cells.

In contrast, BPS promoted a marked decrease in insulin secretion in mouse and in human β-cells, which was already evident at the lowest doses assayed (100 pM, 1 nM). Compared to BPA, the effects of BPS on pancreatic β-cells remain much less well understood. To date, only one study has been published exploring BPS action. This study showed that mouse islets of Langerhans treated with 1 nM and 1 μM BPS exhibited enhanced insulin release compared to controls [37].

The reasons for the discrepancy may include different time treatments, cellular models, and species used. In any case, although few epidemiological studies have been published, they all concluded that BPS is associated with increased risk of type 2 diabetes [38, 39]. A decline in pancreatic β-cell function, as here reported, could be a molecular mechanism underlying this phenomenon.

Our study also looked at gene expression changes caused by the selected EDCs. The analysis, although limited, focused on genes relevant to pancreatic β-cell function and identity. Some of the most prominent changes were observed in BPA-treated murine β-cells with an upregulation of the transcription factors Pdx1, Hnf4α, and Mafa. This correlated quite well with the increase in insulin release as these transcription factors are important for normal glucose sensing and insulin secretion in adult β-cells [40]. These results also agreed with the findings observed in β- cells isolated from BPA-exposed animals at different periods of life [41, 42]. In the case of BPS, we found a consistent downregulation of the glucose transporter gene in both mouse and human β- cells. This was the first evidence that BPS alters mRNA expression of a key player in glucose uptake in pancreatic β-cells. In this regard, we speculate that the biological effects of BPS may be similar in other physiological systems, as this EDC also promoted a downregulation of the sodium glucose transporter (SGLT1) and the glucose transporter 2 (GLUT2) in the duodenum [43].

### DEHP

Consistent with the current literature, we found that DEHP treatment led to impaired insulin secretion. This phenomenon has been reported in other in vitro studies. For example, studies performed on the rat insulinoma cell line INS-1 have shown that cells exposed to DEHP for 24 h at different concentrations in the micromolar range [44, 45] exhibited decreased insulin secretion. A similar response was found in the murine β-cell line Rin 5F [46]. Under in vivo conditions, DEHP has also been shown to negatively affect pancreatic β-cell function and, accordingly, glucose homeostasis [47–51].

Of considerable interest was demonstrating for the first time that DEHP promoted a decline in insulin secretory capacity in human pancreatic β-cells, an effect comparable to that found in murine β-cells, although the response pattern was slightly different. While MIN6 cells manifested marked diminished insulin release in the concentration range from 100 pM to 10 μM, human β- cells exhibited a non-monotonic dose response (NMDR) with reduced insulin secretion at nanomolar doses and an increase at a higher concentration (1 μM). These findings are of relevance as standard risk assessments have as a core assumption that linear extrapolation procedures can be used to predict effects. Of note, NMDRs associated with DEHP exposure have also been observed in a broad range of in vivo and in vitro studies [52]. Some of the outcome variables affected include aromatase activity [53], fetal male serum testosterone, testicular testosterone, and anogenital distance [54]. To the best of our knowledge, this was the first experimental indication that the same behavior could affect the insulin secretory response to glucose.

### PFOS

It is known that PFOS can accumulate in pancreatic tissue [55, 56], affecting pancreatic β-cell function. In particular, a dual effect of PFOS on insulin secretion has been reported that could be attributed to different EDC exposure times or species specificity and sensitivity as discussed below.

Thus, in the murine pancreatic β-cell line TC-6, acute PFOS exposure (5–100 μM) promoted increased insulin secretion in response to low (1.4 mM) [57] and high glucose concentrations (16.7 mM) [50]. On the contrary, 48 h exposure to PFOS (micromolar range 10–200 μM) resulted in decreased insulin secretion together with diminished ATP production, membrane potential, and intracellular calcium concentration [58]. In MIN6 cells and isolated mouse islets, PFOS treatment at 1 and 10 μM doses for 24 h led to reduced ATP production and GSIS, an effect proposed to be mediated by the downregulation of an insulin-related Akt-pathway [59]. These data agreed with the findings reported here. We tested different concentrations of PFOS in the range of 100 pM to 10 μM and found that MIN6 cells treated for 24 h showed a marked decrease in insulin release, an effect that was more significant at 100 pM, 100 nM, and 10 μM concentrations. Importantly, we found that this was correlated with a reduced gene expression of Glut2. Therefore, our data not only confirmed the disruptive effect of PFOS on pancreatic β-cell function but also provided new evidence highlighting that this may occur not only in the micromolar range but also at lower PFOS doses. This is extremely relevant since lower doses seem to be more realistic in terms of human exposure.

Of note, we also found that human pancreatic β-cells treated with 10 and 100 nM PFOS for 72 h showed enhanced insulin secretory activity and content, a response similar to the one revealed in human population studies. Many epidemiological investigations have demonstrated that plasma PFOS concentrations are positively correlated with elevated insulin levels, augmented pancreatic β-cell activity, and insulin resistance in adults [60–63] as well as in overweight children [64]. In vivo animal studies have pointed in the same direction and shown that developmental exposure to PFOS may lead to hyperinsulinemia and insulin resistance in offspring when reaching adulthood [65].

### CdCl_2_

It has been reported that pancreatic β-cells are particularly susceptible to the accumulation of CdCl_2_ in a time- and dose-dependent manner. Furthermore, it has been pointed out that CdCl_2_ may compete with Zn^2+^ for some of its transporters and binding proteins [66]. This could be quite relevant considering that Zn^2+^ plays a key role in the correct storage and release of insulin [67]. It is then reasonable to speculate that the potential harmful effects of this metal on β-cells will depend to a large extent on both factors. A study carried out on the MIN6 β-cell line showed that in vitro cellular treatment with CdCl_2_ (0.5 and 1 μM) for 48 h decreased insulin secretion in a concentration-dependent manner with no effect at the lowest dose assayed [66]. In another study, MIN6 cells were exposed to CdCl_2_ (1–5 μM) for 24 h, and diminished insulin release was found in response to 2 μM CdCl_2_ concentration or above [68], while in Rin-m5F cells [69] higher doses (5 and 10 μM) were required to achieve the same effect. Our data indicated that treatment of MIN6 cells with CdCl_2_ in a range of doses from 100 pM to 1 μM for 24 h did not exert any effect on GSIS. Given the above, we hypothesized that longer exposure time and higher doses are required for CdCl_2_ to have an impact on β-cell function. Surprisingly, in human β-cells we found that 10 nM CdCl_2_ treatment for 72 h led to a modest increase in insulin release although no effect was observed at lower (1 nM) or higher (100 nM, 1 μM) doses. This was in accordance with an increase in Ca^2+^ currents at 10 nM, which could explain this slight increase in insulin secretion. Whether or not this has a major impact on pancreatic β-cell physiology should be further explored.

In vivo studies also suggested that CdCl_2_ effects on pancreatic β-cell activity and glucose homeostasis may vary, presumably depending on time, doses of exposure, and route of administration. For example, rats orally exposed to CdCl_2_ for 1 month manifested hyperinsulinemia and insulin resistance [70, 71]. Longer exposures to CdCl_2_ (12 w) led to diminished insulin secretion in isolated islets [72], while rats intraperitoneally injected with CdCl_2_ for 24 w manifested decreased serum insulin levels and liver insulin receptor expression, but no effects on glucose levels or insulin tolerance were detected [73].

### DDE

DDE is the major metabolite of the persistent organochloride pesticide DDT. Although there is some epidemiological evidence indicating a positive association between DDE levels and diabetes incidence [74, 75], whether this results from a direct action of DDE on pancreatic β-cell function remains unknown. To date, very few studies have explored this plausible underlying mechanism. Previous studies have reported that basal insulin levels decreased in β-TC-6 and INS-1 cells exposed to 10 μM DDE, but none analyzed the effects of DDE on the insulin secretory response to stimulatory glucose concentrations. In our experiments, we did not find any effect of DDE on insulin secretion or content, either in mouse β-cells or in human cells, at least under the experimental conditions assayed. In any case, more investigation is needed in order to decipher DDE impact on β-cell physiology.

In summary, as previously discussed, we provided evidence that the selected EDCs impaired a number of metabolic endpoints in pancreatic β-cells. It is worth mentioning that these endpoints can be considered important molecular key events (KEs) in an adverse outcome pathway (AOP) framework for diabetes. These KEs include: i) impaired expression of genes encoding for β-cell function and identity, ii) decreased K^+^ currents, iii) increased Ca^2+^ currents, and iii) altered insulin secretion and content. This is of considerable relevance as KEs are essential biological events connecting a molecular initiate event (MIE) with a final adverse outcome (AO) like insulin resistance or hyperglycemia.

## 4. Conclusions

Our results showed that both pancreatic β-cell models, MIN6 and EndoC-βH1, were responsive and sensitive to most of the EDCs tested and that the responses were in general agreement with those revealed in other in vivo and in vitro studies found in the literature. In addition, we revealed for the first time the impact of relevant EDCs on key molecular aspects of pancreatic β- cell physiology, such as electrical activity and insulin release in a human cell-based model. As such, this system may shed light on the molecular mechanisms underlying the EDC mode of action and improve the risk assessment framework for MDC in relation to human β-cell dysfunction. Overall, the work presented here points to the possibility of using the pancreatic β- cell lines assayed as sensitive screening tools for the identification of potential diabetogenic environmental pollutants.

## 5. Material and methods

### 5.1. Chemicals

Bisphenol A (BPA, Cat No 239658), bisphenol S (BPS, Cat No 103039), bisphenol F (BPF, Cat No B47006), Di(2-ethylhexyl) phthalate (DEHP, Cat No 36735), perfluorooctanesulfonic acid (PFOS, Cat No 77283), cadmium chloride (CdCl2, Cat No 202908), and 1,1-Dichloro-2,2-bis(4-chlorophenyl)ethene, 4,4′-DDE (4,4′-DDE, Cat No 35487), and IBMX (Cat No I-5879), were purchased from Sigma-Aldrich (Saint Louis, MO, USA). Diazoxide (Cat no 0964) was purchased from Tocris Cookson (Bristol, UK). All chemicals were dissolved in dimethyl sulfoxide (DMSO) to prepare the stock solution, except for CdCl_2_, which was dissolved in water. The amount of DMSO for exposure studies was maintained at less than 0.03%.

### 5.2. Cell culture

The MIN-6 cell line was kindly provided by Dr. Jun-Ichi Miyazaki (Osaka University, Osaka, Japan). MIN6 cells were grown in optimized DMEM (AddexBioTechnologies) supplemented with 10% FBS (HyClone, GE Healthcare Life Sciences), 50 μM 2-β-mercaptoethanol (Gibco, UK), 100 units/mL penicillin and 100 μg/mL streptomycin (Thermo Fisher Scientific, Waltham, MA, USA). For EDC treatment, DMEM without phenol red (Sigma-Aldrich, Saint Louis, MO, USA) was used. It was supplemented with 1.5 g/l sodium bicarbonate, 1 mM sodium pyruvate (Gibco, UK), 50 μM 2-β-mercaptoethanol (Gibco, UK), 100 units/mL penicillin, 100 μg/mL streptomycin (Thermo Fischer Scientific), and 10% charcoal dextran-treated FBS (HyClone, GE Healthcare Life Sciences).

EndoC-βH1 cells (Univercell biosolutions, Paris, France) were cultured and passaged as previously described [76]. Briefly, cells were cultured in DMEM containing 5.6 mM glucose, 2% BSA fraction V fatty acid free (Sigma-Aldrich, Saint Louis, MO, USA), 50 μM 2-β- mercaptoethanol (Gibco, UK), 10 mM nicotinamide, 5.5 μg/ml human transferrin, 6.7 ng/ml sodium selenite (Sigma-Aldrich, Saint Louis, MO, USA), 100 units/mL penicillin and 100 μg/mL streptomycin (Thermo Fischer Scientific, Waltham, MA, USA). Cells were seeded at a density of 70–80*10^3^ cells/cm^2^ on ECM (1%) and fibronectin (2 μg/ml) (Sigma-Aldrich, Saint Louis, MO, USA) -coated plates, and cultured at 37 °C in 5% CO_2_. For treatment with EDCs, DMEM was replaced with DMEM without phenol red (Sigma-Aldrich, Saint Louis, MO, USA) and BSA was replaced by 2% charcoal dextran-treated FBS (HyClone, GE Healthcare Life Sciences).

Both cell lines were incubated at 37 °C and 5% CO_2_. Cells were discarded after passage 32 for MIN6 and passage 72 for EndoC-βH1.

### 5.3. Cell viability

MIN6 and EndoC-βH1 cells were seeded at densities of 35,000 and 20,000 cells per well, respectively, in dark 96-well plates (Corning Incorporated, Kennebunk, USA) 72 h before the treatment. Then, the medium was replaced, and cells were treated with vehicle or different concentrations of the EDCs tested for 24, 48, 72 h or 7 d as indicated in the figure legends.

The medium was replaced every 24 h. At the end of the incubation time cell viability was assessed using a combination of three indicator dyes: RZ (Thermo Fisher Scientific, Waltham, MA, USA), neutral red (Sigma-Aldrich, Saint Louis, MO, USA), and 5-carboxyfluorescein diacetate acetoxymethyl ester (CFDA-AM; Thermo Fisher Scientific, Waltham, MA, USA). Cells were washed with phosphate-buffered saline (PBS) and incubated for 40 min with a solution of RZ (5% v/v) and CFDA-AM (4 μM) prepared in serum-free DMEM. After incubation, fluorescence (RZ: excitation 530–570 nm, emission 590–620 nm; CFDA-AM: excitation 485 nm, emission 520 nm) was measured using a fluorescence plate reader (POLARstar Omega, BMG Labtech). After removing the RZ and CFDA-AM dye, cells were rinsed twice with PBS and incubated for 2 h with neutral red solution (0.005% w/v). Accumulated neutral red was extracted from cells by a lysis solution (1% glacial acetic acid in 50% ethanol). NRU was quantified by the measurement of absorbance at 540 and 690 nm (background) using a microplate reader (Biotek EON). An amount of 3 or 10% DMSO was used as positive control for cellular damage in MIN6 and EndoC-βH1, respectively. The results are expressed as percentages (%) of the readings in the control wells.

### 5.4. Insulin secretion and content

EndoC-βH1cells were seeded onto coated 24-well plates at a density of 200*10^3^ cells per well for 72 h before EDC treatment. Then, cells were exposed to different concentrations of test EDCs for 48, 72 h or 7 d (exposure time was optimized for each compound). Prior to the insulin secretion experiment, cells were washed twice and preincubated for 2 h in a modified Krebs–Ringer medium containing 120 mM NaCl, 5.4 mM KCl, 1.2 mM KH_2_PO_4_, 1.2 mM MgSO_4_, 20 mM HEPES, 2.4 mM CaCl_2_ and 0.1% BSA, pH 7.4 at 37 °C. Cells were then incubated with Krebs–Ringer medium at 2.8 mM glucose for 40 min, followed by 20 mM glucose for 40 min. 3-Isobutyl-1- methylxanthine (IBMX) (0.5 mM) was used as a positive control and diazoxide (DZX) (0.2 mM) as a negative control. Supernatants were collected as basal and stimulated insulin secretions, respectively, centrifuged at 700 G and 4 °C for 5 min, and immediately frozen for later analysis. For insulin content cells were lysed using TETG buffer supplemented with Complete™ Mini Anti-protease (Sigma-Aldrich, Saint Louis, MO, USA) (prepared following the manufacturer instructions). Lysate was centrifuged at 700 G and 4 °C for 5 min and supernatants were immediately frozen. Insulin secretion and content were measured by ELISA according to manufacturer’s instructions using the Human Insulin Kit (Mercodia, Sweden). Protein measurement was carried out using Pierce BCA kit (Thermo Fisher). Insulin secretion and content were normalized by protein content and expressed as percentage of control 2.8 mM glucose.

MIN6 cells were seeded in 24-well plates at a density of 150*10^3^ cells per well for 72 h before EDC treatment. Cells were preincubated with Krebs–Ringer medium (0 mM glucose) for 2 h and then incubated with Krebs in 1.67 mM glucose for 30 min, followed by 16.7 mM glucose for 30 min. Streptozotocin (STZ) (0.25 mM) was used as negative control. For insulin content measurement, cells were washed with PBS, lysed in ice-cold acid ethanol (75% ethanol, 1.5% HCl) and incubated overnight at 4 °C. Insulin secretion and content were measured by ELISA (Mercodia, Sweden). Protein measurement was carried out using Bradford (Sigma-Aldrich). Insulin secretion and content were normalized by protein content and expressed as percentage of control 1.67 mM glucose. For simplicity, only significant differences due to EDC treatment between low or high-glucose and vehicle-treated cells are denoted.

### 5.5. RNA extraction and RT-qPCR

Pancreatic cells were seeded in 24-well plates at a density of 150*10^3^ cells/well for MIN6 cells and 200*10^3^ cells/well for EndoC-βH1 cells. After EDC treatment, RNA was extracted using a commercial kit (RNeasy Micro kit, Qiagen, Hilden, Germany) according to the manufacturer’s instructions. RNA (1 μg) was reverse transcribed using the High Capacity cDNA Reverse Transcription kit (Applied Biosystems, Foster City, CA, USA). Quantitative PCR assays were performed using the CFX96 Real Time System (Bio-Rad, Hercules, CA, USA). Amplification reactions were carried out in medium containing a 200 nM concentration of each primer, 1 μl of cDNA, and 1x IQ SYBR Green Supermix (Bio-Rad). Primers were designed between exons to avoid genomic cross-reaction. Samples were subjected to the following conditions: 30 s at 95 °C, 45 cycles (5 s at 95 °C, 5 s at 60 °C, and 10 s at 72 °C) and a melting curve of 65–95 °C. The resulting values were analyzed with the CFX96 Real-Time System (Bio-Rad Laboratories, Hercules, CA, USA) and were expressed relative to the control values (2^−ΔΔCT^). All measurements were performed in duplicate and normalized against the geometric mean of the housekeeping genes Actb and Hprt. The primers used herein are listed in Supplemental Table 3.

### 5.6. Electrophysiological Recordings: K^+^ and Ca^2+^ currents

EndoC-βH1 cells were plated on ECM–fibronectin pre-coated slip slides (10 mm) at a density of 150*10^3^ cells per slide. After at least 24 h cells were exposed to EDCs at different concentrations as indicated in the figure legend. For the patch-clamp recordings of voltage-gated K^+^ and Ca^2+^ currents the standard whole-cell patch-clamp was used, as previously described [25]. Data were obtained using an Axopatch 200B amplifier (Axon Instruments Co., Union City, CA, USA). Patch pipettes were pulled from borosilicate capillaries (Sutter Instruments Co., Novato, CA, USA) using a flaming/brown micropipette puller P-97 (Sutter Instruments Co.Novato, CA, USA) and heat polished at the tip using an MF-830 microforge (Narishige, Japan). The bath solution contained: 135 mM NaCl, 5 mM KCl, 2.5 mM CaCl_2_, 1.1 mM MgCl_2_, 10 mM HEPES, and 5 mM glucose (pH: 7.4 with NaOH). Various pipette-filling solutions were used. For recordings of voltage-gated K^+^ currents, the pipette was filled with: 120 mM KCl, 1 mM MgCl_2_, 1 mM CaCl_2_, 3 mM MgATP, 10 mM EGTA, and 10 mM HEPES (pH: 7.15 with KOH). A similar medium was used for the Ca^2+^ current measurements except that KCl was equimolarly replaced by CsCl and pH adjusted with CsOH (pH: 7.15). K^+^ currents were recorded in response to depolarizing voltage pulses of –60 mV to +80 mV from a holding potential of –70 mV. Ca^2+^ currents were recorded in response to depolarizing voltage pulses of –60 mV to +50 mV from a holding potential of –70 mV. For K^+^ and Ca^2+^ currents density quantification, K^+^ and Ca^2+^ currents (in pA) were normalized to the cell capacitance (in pF).

After filling the pipette with the pipette solution, the pipette resistance was 3–5 MΩ. A tight seal (>1 GΩ) was established between the cell membrane and the tip of the pipette by gentle suction. The series resistance of the pipette usually increased to 6–15 MΩ after moving to the whole cell. Series resistance compensation was used (up to 70%) to keep the voltage error below 5 mV during current flow. Finally, data were filtered (2 KHz) and digitized (10 KHz) using a Digidata 1550B1 (Molecular Devices, CA, USA).

### 5.7 Statistical analysis

The GraphPad Prism 7.0 software (GraphPad Software, Inc., CA, USA) was used for all statistical analyses. Data are expressed as the mean ± SEM. To assess differences between groups, two-way analysis of variance (ANOVA) followed by post hoc Tukey test, one-way ANOVA followed by Dunnett’s test, or Student’s t-test was used when appropriate. When data did not pass the parametric test, Kruskal–Wallis followed by Dunn’s multiple comparison post hoc test was used. Statistical significance was set at p < 0.05 for all the analyses. The statistical tests used in each experiment are specified in each figure legend.

## Supplementary Materials

Figure S1; Figure S2, Figure S3, Figure S4, Table S1, Table S2, Table S3.

## Author Contributions

Conceptualization, RAA, HF and PAM; formal analysis, RAA, HF, SS, TBB and PAM; investigation, RAA, HF, SS, TBB and PAM; resources, PAM; writing—original draft preparation, PAM; writing—review and editing, RAA, HF, SS, TBB and PAM.; visualization, RAA, HF, SS, TBB and PAM; supervision, PAM; funding acquisition, PAM. All authors have read and agreed to the published version of the manuscript.

## Acknowledgments

The authors thank M. L. Navarro and S. Ramon (IDiBE, Universidad Miguel Hernández) for their excellent technical assistance.

