## Supplementary material for "Screening of Relevant Metabolism-Disrupting Chemicals on Pancreatic β-Cells: Evaluation of Murine and Human in Vitro Models"

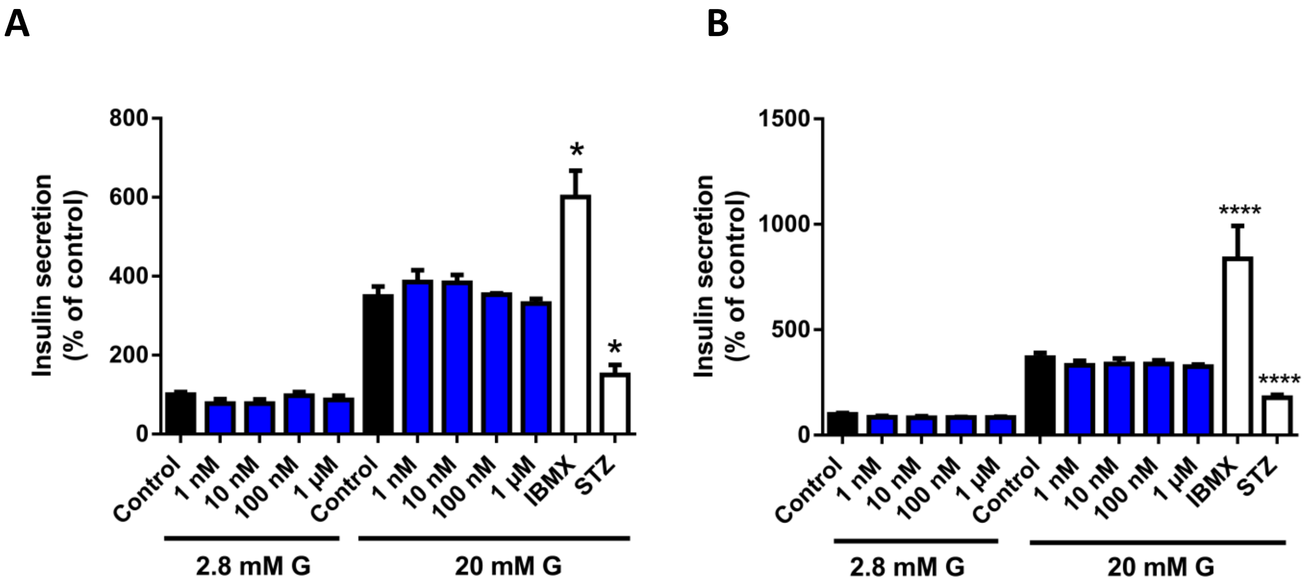


**Supplemental Figure 1.** Glucose-stimulated insulin secretion (GSIS) in EndoC-βH1 cells treated with BPA (1 nM–1 µM) for 24 h (A) or 48 h (B). n = two independent experiments (A), n = three independent experiments (B). * vs. Control 20 mM G; *p < 0.05, ****p < 0.0001, one-way ANOVA followed by Dunnet's post hoc test. All data are expressed as mean ± SEM.

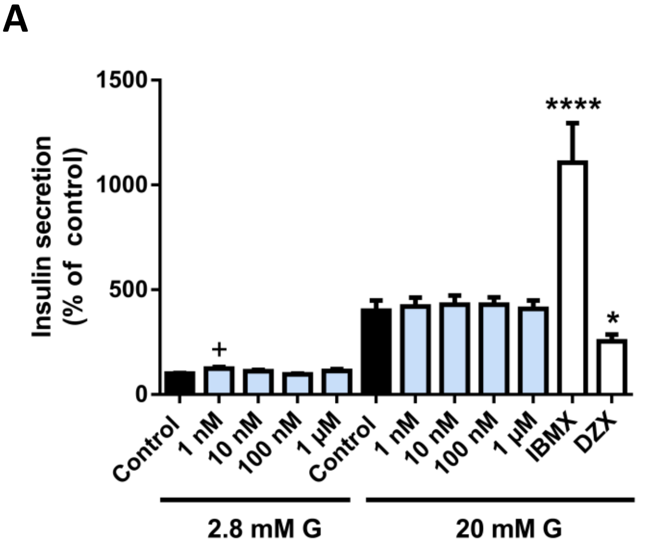


B

| 24 h | Resazurin | Neutral red | CFDA-AM |
|---|---|---|---|
| Control | 100.0 ± 0.5 | 100.0 ± 1.6 | 100.0 ± 2.3 |
| BPF 1 nM | 98.4 ± 0.6 | 102.3 ± 1.9 | 98.4 ± 2.1 |
| BPF 10 nM | 96.1 ± 1.0 * | 104.6 ± 1.9 | 94.9 ± 2.1 |
| BPF 100 nM | 94.0 ± 1.3 *** | 107.1 ± 2.0 | 90.4 ± 2.3 ** |
| BPF 1 µM | 91.2 ± 1.2 **** | 109.5 ± 3.0 ** | 92.5 ± 2.5 * |
| 48 h | Resazurin | Neutral red | CFDA-AM |
| Control | 100.0 ± 0.5 | 100.0 ± 1.7 | 100.0 ± 1.6 |
| BPF 1 nM | 96.2 ± 0.8 | 103.1 ± 1.3 | 95.5 ± 1.8 |
| BPF 10 nM | 93.2 ± 0.4 *** | 104.2 ± 2.1 | 99.8 ± 2.0 |
| BPF 100 nM | 87.9 ± 1.2 **** | 106.7 ± 1.6 * | 94.0 ± 1.6 |
| BPF 1 µM | 90.0 ± 1.2 **** | 106.0 ± 0.9 * | 94.2 ± 2.6 |
| 72 h | Resazurin | Neutral red | CFDA-AM |
| Control | 100.0 ± 0.7 | 100.0 ± 1.2 | 100.0 ± 1.6 |
| BPF 1 nM | 99.8 ± 0.7 | 106.0 ± 1.6 * | 94.7 ± 2.9 |
| BPF 10 nM | 98.8 ± 1.2 | 108.5 ± 1.8 *** | 98.2 ± 3.1 |
| BPF 100 nM | 96.6 ± 1.0 | 106.2 ± 1.7 * | 94.6 ± 2.5 |
| BPF 1 µM | 94.5 ± 1.2 *** | 105.4 ± 1.1 * | 96.3 ± 2.2 |

**Supplemental Figure 2.** (A) GSIS in EndoC-βH1 cells treated with BPF (1 nM–1 µM) for 72 h. n = three independent experiments. + vs. Control 2.8 mM G and * vs. Control 20 mM G; +p < 0.05, *p < 0.05, ****p < 0.0001, one-way ANOVA followed by Dunnet's post hoc test. (B) Viability of EndoC-βH1 cells treated for 72 h with different BPF concentrations (1 nM-1 µM) as evaluated by RZ, NR and CFDA-AM assays. n = four independent experiments. * vs. Control; *p < 0.05, **p < 0.01, ***p < 0.001 and

****p < 0.0001, one-way ANOVA followed by Dunnet's post hoc test or Kruskal-Wallis followed by Dunn's post hoc test. All data are expressed as mean ± SEM.

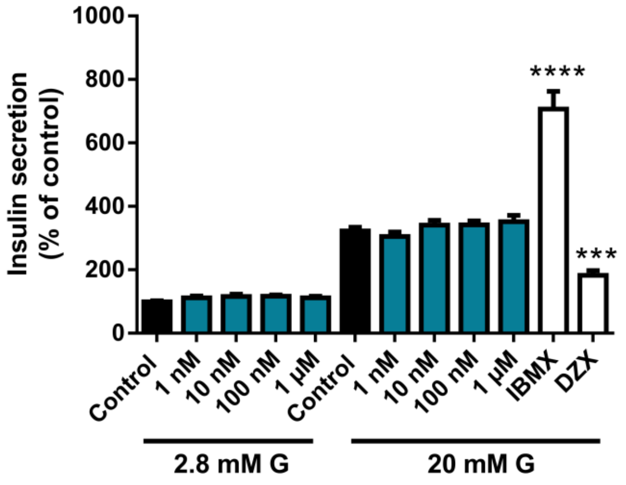


**Supplemental Figure 3.** GSIS in EndoC-βH1 cells treated with DEHP (1 nM–1 µM) for 72 h. n = six independent experiments. * vs. Control 20 mM G; ***p < 0.001, ****p < 0.0001, one-way ANOVA followed by Dunnet's post hoc test. All data are expressed as mean ± SEM.

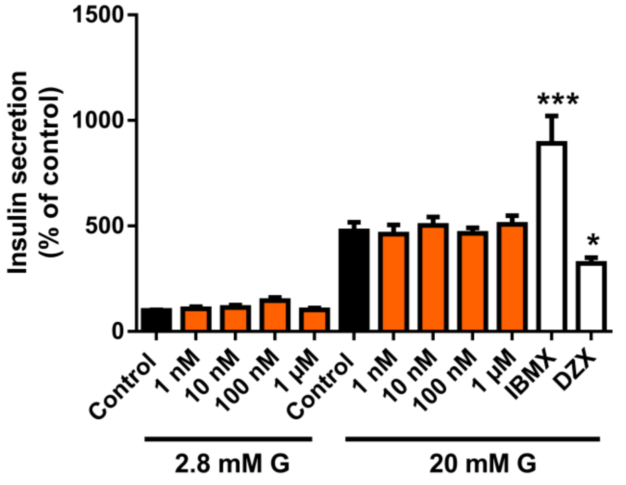


**Supplemental Figure 4.** GSIS in EndoC-βH1 cells treated with DDE (1 nM–1 µM) for 72 h. n = three independent experiments. * vs. Control 20 mM G; *p < 0.05, ***p < 0.001, ANOVA one way followed by Dunnet's post hoc test. All data are expressed as mean ± SEM.

| MIN6 | | | | | | | |
|---|---|---|---|---|---|---|---|
| **48 h** | **Resazurin** | **Neutral red** | **CFDA-AM** | **48 h** | **Resazurin** | **Neutral red** | **CFDA-AM** |
| **Control** | 100.0 ± 0.9 | 100.0 ± 0.9 | 100.0 ± 0.8 | **Control** | 100.0 ± 1.9 | 100.0 ± 0.8 | 100.0 ± 0.7 |
| **BPA 100 pM** | 93.5 ± 1.4 * | 99.9 ± 1.1 | 96.0 ± 1.0 | **BPS 100 pM** | 90.6 ± 2.8 * | 93.3 ± 1.5 *** | 96.5 ± 1.7 |
| **BPA 1 nM** | 86.4 ± 2.3 **** | 100.5 ± 1.3 | 90.2 ± 1.6 **** | **BPS 1 nM** | 89.0 ± 3.3 * | 96.8 ± 1.3 | 96.6 ± 1.2 |
| **BPA 10 nM** | 87.7 ± 1.6 **** | 93.3 ± 1.8 | 92.4 ± 1.2 *** | **BPS 10 nM** | 89.2 ± 3.1 * | 96.3 ± 1.0 | 95.1 ± 1.3 * |
| **BPA 100 nM** | 89.7 ± 1.5 *** | 100.7 ± 1.3 | 94.4 ± 1.4 * | **BPS 100 nM** | 89.6 ± 2.9 * | 97.5 ± 1.1 | 94.9 ± 1.2 * |
| **BPA 1 µM** | 85.6 ± 2.2 **** | 85.1 ± 3.2 * | 88.1 ± 1.2 **** | **BPS 1 µM** | 87.0 ± 2.6 ** | 96.6 ± 1.0 | 92.9 ± 1.1 *** |
| **BPA 10 µM** | 87.4 ± 3.4 ** | 100.1 ± 1.6 | 92.0 ± 1.5 **** | **BPS 10 µM** | 84.1 ± 2.3 **** | 89.9 ± 1.4 **** | 93.0 ± 1.2 *** |
| **72 h** | **Resazurin** | **Neutral red** | **CFDA-AM** | **72 h** | **Resazurin** | **Neutral red** | **CFDA-AM** |
| **Control** | 100.0 ± 1.2 | 100.0 ± 1.2 | 100.0 ± 0.7 | **Control** | 100.0 ± 2.0 | 100.0 ± 0.7 | 100.0 ± 1.1 |
| **BPA 100 pM** | 89.3 ± 2.8 * | 92.4 ± 3.7 | 94.4 ± 1.6 * | **BPS 100 pM** | 97.3 ± 2.3 | 100.5 ± 0.9 | 98.1 ± 1.2 |
| **BPA 1 nM** | 94.1 ± 2.0 | 103.9 ± 2.0 | 98.2 ± 1.2 | **BPS 1 nM** | 92.6 ± 2.3 | 101.3 ± 0.9 | 94.6 ± 1.3* |
| **BPA 10 nM** | 78.2 ± 1.9 **** | 85.3 ± 2.3 ** | 88.4 ± 1.2 **** | **BPS 10 nM** | 90.9 ± 2.4 * | 99.9 ± 1.1 | 94.3 ± 1.4** |
| **BPA 100 nM** | 103.8 ± 2.0 | 116.0 ± 2.5 ** | 102.3 ± 0.9 | **BPS 100 nM** | 92.2 ± 2.1 | 102.9 ± 1.0 | 94.8 ± 1.3* |
| **BPA 1 µM** | 96.2 ± 2.7 | 107.5 ± 3.6 | 96.3 ± 1.5 | **BPS 1 µM** | 94.3 ± 2.1 | 102.5 ± 1.1 | 95.2 ± 1.2* |
| **BPA 10 µM** | 82.8 ± 3.1 **** | 83.9 ± 3.5 ** | 88.9 ± 1.8 **** | **BPS 10 µM** | 93.7 ± 2.1 | 99.9 ± 1.7 | 95.2 ± 1.3* |
| **48 h** | **Resazurin** | **Neutral red** | **CFDA-AM** | **48 h** | **Resazurin** | **Neutral red** | **CFDA-AM** |
| **Control** | 100.0 ± 1.1 | 100.0 ± 1.0 | 100.0 ± 0.9 | **Control** | 100.0 ± 3.0 | 100.0 ± 0.7 | 100.0 ± 4.2 |
| **PFOS 100 pM** | 91.9 ± 1.9 ** | 93.4 ± 1.9 * | 95.6 ± 1.5 * | **$CdCl_2$ 100 pM** | 92.6 ± 1.6 | 87.1 ± 3.8 | 91.5 ± 3.5 |
| **PFOS 1 nM** | 94.4 ± 1.8 * | 93.5 ± 2.5 ** | 99.7 ± 2.0 | **$CdCl_2$ 1 nM** | 83.3 ± 3.5 ** | 71.1 ± 2.1 **** | 86.7 ± 2.7 |
| **PFOS 10 nM** | 89.2 ± 1.2 **** | 83.9 ± 2.1 **** | 94.5 ± 0.9 ** | **$CdCl_2$ 10 nM** | 83.9 ± 3.2 * | 83.3 ± 3.2 | 86.1 ± 3.0 |
| **PFOS 100 nM** | 92.9 ± 1.5 ** | 96.8 ± 1.9 | 97.4 ± 1.0 | **$CdCl_2$ 100 nM** | 80.2 ± 2.5 ** | 72.1 ± 3.3 *** | 78.2 ± 5.0 * |
| **PFOS 1 µM** | 83.7 ± 2.5 **** | 82.7 ± 1.7 **** | 86.9 ± 1.8 **** | **$CdCl_2$ 1 µM** | 63.4 ± 2.4 **** | 56.2 ± 2.7 **** | 57.9 ± 3.5 **** |
| **PFOS 10 µM** | 95.0 ± 1.1 | 92.7 ± 1.1 * | 94.8 ± 1.4 ** | **$CdCl_2$ 10 µM** | 9.7 ± 2.3 **** | 2.2 ± 0.9 **** | 17.1 ± 3.6 **** |
| **72 h** | **Resazurin** | **Neutral red** | **CFDA-AM** | **72 h** | **Resazurin** | **Neutral red** | **CFDA-AM** |
| **Control** | 100.0 ± 1.7 | 100.0 ± 2.3 | 100.0 ± 0.9 | **Control** | 100.0 ± 1.8 | 100.0 ± 0.6 | 100.0 ± 1.3 |
| **PFOS 100 pM** | 97.6 ± 2.3 | 100.9 ± 2.2 | 97.3 ± 1.2 | **$CdCl_2$ 100 pM** | 92.2 ± 2.0 | 95.9 ± 0.8 * | 95.3 ± 1.2 |
| **PFOS 1 nM** | 95.6 ± 2.3 | 101.3 ± 2.2 | 96.2 ± 1.2 | **$CdCl_2$ 1 nM** | 88.5 ± 2.3 ** | 92.1 ± 0.8 **** | 94.2 ± 1.5 |
| **PFOS 10 nM** | 94.2 ± 2.5 | 97.3 ± 2.4 | 96.8 ± 1.2 | **$CdCl_2$ 10 nM** | 89.5 ± 2.2 * | 91.9 ± 0.8 **** | 94.0 ± 1.4 |
| **PFOS 100 nM** | 97.3 ± 2.6 | 101.7 ± 2.8 | 96.6 ± 1.3 | **$CdCl_2$ 100 nM** | 91.1 ± 2.3 | 92.3 ± 1.2 **** | 94.1 ± 1.5 |
| **PFOS 1 µM** | 97.3 ± 2.1 | 99.8 ± 2.3 | 96.8 ± 1.1 | **$CdCl_2$ 1 µM** | 86.9 ± 2.7 ** | 90.1 ± 1.4 **** | 92.4 ± 1.6 ** |
| **PFOS 10 µM** | 102.3± 2.3 | 101.0 ± 2.8 | 98.5 ± 1.1 | **$CdCl_2$ 10 µM** | 0 ± 0.3 **** | 53.2 ± 1.4 **** | 10.8 ± 0.4 **** |
| **48 h** | **Resazurin** | **Neutral red** | **CFDA-AM** | **48 h** | **Resazurin** | **Neutral red** | **CFDA-AM** |
| **Control** | 100.0 ± 1.6 | 100.0 ± 1.0 | 100.0 ± 1.0 | **Control** | 100.0 ± 0.9 | 100.0 ± 0.8 | 100.0 ± 1.0 |
| **DEHP 100 pM** | 95.8 ± 1.9 | 99.0 ± 1.4 | 96.7 ± 1.5 | **DDE 100 pM** | 95.6 ± 1.3 | 102.9 ± 1.5 | 97.0 ± 1.5 |
| **DEHP 1 nM** | 94.1 ± 2.0 | 97.8 ± 1.0 | 96.7 ± 1.3 | **DDE 1 nM** | 95.6 ± 1.5 | 100.9 ± 2.3 | 95.9 ± 1.2 |
| **DEHP 10 nM** | 92.9 ± 2.2 | 94.1 ± 1.3 ** | 95.5 ± 1.7 | **DDE 10 nM** | 85.4 ± 1.8 **** | 89.1 ± 1.9 ** | 93.5 ± 1.6 ** |
| **DEHP 100 nM** | 94.1 ± 2.2 | 99.6 ± 1.4 | 96.0 ± 1.4 | **DDE 100 nM** | 94.4 ± 1.3 * | 101.2 ± 1.8 | 99.9 ± 1.4 |
| **DEHP 1 µM** | 91.3 ± 2.5 * | 93.5 ± 1.2 ** | 94.7 ± 1.4 * | **DDE 1 µM** | 91.1 ± 1.4 *** | 95.2 ± 1.5 | 96.1 ± 1.3 |
| **DEHP 10 µM** | 92.5 ± 3.5 | 79.5 ± 1.7 **** | 83.2 ± 3.2 **** | **DDE 10 µM** | 77.4 ± 4.1 **** | 79.5 ± 4.5 **** | 87.6 ± 2.2 *** |
| **72 h** | **Resazurin** | **Neutral red** | **CFDA-AM** | **72 h** | **Resazurin** | **Neutral red** | **CFDA-AM** |
| **Control** | 100.0 ± 1.6 | 100.0 ± 1.7 | 100.0 ± 1.4 | **Control** | 100.0 ± 1.7 | 100.0 ± 1.0 | 100.0 ± 1.1 |
| **DEHP 100 pM** | 94.3 ± 1.9 | 96.2 ± 1.8 | 93.6 ± 0.9 ** | **DDE 100 pM** | 99.5 ± 3.7 | 111.4 ± 2.7 * | 99.1 ± 1.6 |
| **DEHP 1 nM** | 95.4 ± 1.6 | 99.0 ± 2.0 | 95.3 ± 1.0 * | **DDE 1 nM** | 94.4 ± 4.1 | 106.8 ± 2.8 | 96.6 ± 2.0 |
| **DEHP 10 nM** | 93.3 ± 1.9 * | 96.8 ± 1.4 | 95.2 ± 1.0 * | **DDE 10 nM** | 96.5 ± 4.7 | 113.5 ± 3.0 ** | 97.9 ± 2.3 |
| **DEHP 100 nM** | 95.3 ± 1.7 | 99.2 ± 2.1 | 96.8 ± 0.9 | **DDE 100 nM** | 94.5 ± 5.0 | 113.9 ± 3.2 ** | 96.9 ± 2.4 |
| **DEHP 1 µM** | 88.6 ± 2.2 *** | 90.7 ± 1.9 ** | 94.6 ± 1.0 * | **DDE 1 µM** | 93.5 ± 4.5 | 109.1 ± 2.6 | 95.1 ± 2.1 |
| **DEHP 10 µM** | 59.1 ± 3.7 **** | 47.1 ± 2.9 **** | 65.9 ± 3.3 **** | **DDE 10 µM** | 40.7 ± 2.4 * | 82.4 ± 6.2 | 69.9 ± 5.1 * |

**Supplemental Table 1.** Viability of MIN6 cells treated for 48 or 72 h with different BPA, BPS, $CdCl_2$, PFOS, DDE or DEHP concentrations (100 pM–10 µM) as evaluated by RZ, NR and CFDA-AM assays. n= at least three independent experiments. * vs. Control; *$p < 0.05$, **$p < 0.01$, ***$p < 0.001$ and ****$p < 0.0001$, one-way ANOVA followed by Dunnet’s post hoc test or Kruskal-Wallis followed by Dunn’s post hoc test. All data are expressed as mean ± SEM.

| EndoC-βH1 | | | | | | | |
|---|---|---|---|---|---|---|---|
| **24 h** | **Resazurin** | **Neutral red** | **CFDA-AM** | **24 h** | **Resazurin** | **Neutral red** | **CFDA-AM** |
| **Control** | 100.0 ± 0.5 | 100.0 ± 2.0 | 100.0 ± 0.8 | **Control** | 100.0 ± 1.0 | 100.0 ± 1.2 | 100.0 ± 0.7 |
| **BPA 1 nM** | 100.0 ± 1.5 | 103.5 ± 1.8 | 98.3 ± 2.0 | **BPS 1 nM** | 103.8 ± 1.4 | 104.4 ± 1.8 | 102.8 ± 1.3 |
| **BPA 10 nM** | 99.1 ± 1.4 | 104.3 ± 1.5 | 99.0 ± 1.3 | **BPS 10 nM** | 105.4 ± 1.2 * | 107.5 ± 2.0 * | 104.4 ± 1.3 |
| **BPA 100 nM** | 98.3 ± 1.3 | 104.2 ± 2.2 | 97.9 ± 1.1 | **BPS 100 nM** | 102.7 ± 1.6 | 101.3 ± 3.2 | 104.7 ± 2.4 |
| **BPA 1 μM** | 96.2 ± 1.4 | 103.5 ± 2.8 | 96.2 ± 1.3 | **BPS 1 μM** | 95.9 ± 1.9 | 98.4 ± 1.2 | 95.7 ± 1.6 |
| **48 h** | **Resazurin** | **Neutral red** | **CFDA-AM** | **72 h** | **Resazurin** | **Neutral red** | **CFDA-AM** |
| **Control** | 100.0 ± 0.9 | 100.0 ± 1.8 | 100.0 ± 1.2 | **Control** | 100.0 ± 0.9 | 100.0 ± 1.5 | 100.0 ± 0.7 |
| **BPA 1 nM** | 100.4 ± 0.5 | 106.9 ± 1.2 | 99.3 ± 0.8 | **BPS 1 nM** | 100.7 ± 1.5 | 105.0 ± 2.5 | 98.7 ± 2.1 |
| **BPA 10 nM** | 98.3 ± 1.0 | 105.6 ± 2.0 | 97.1 ± 1.1 | **BPS 10 nM** | 99.2 ± 2.3 | 103.3 ± 2.4 | 99.4 ± 1.9 |
| **BPA 100 nM** | 96.1 ± 1.4 | 104.6 ± 2.3 | 94.7 ± 1.5 * | **BPS 100 nM** | 104.3 ± 1.8 | 110.9 ± 2.9 ** | 100.4 ± 2.5 |
| **BPA 1 μM** | 94.9 ± 1.5 ** | 98.1 ± 3.4 | 93.1 ± 1.5 *** | **BPS 1 μM** | 97.5 ± 1.8 | 97.0 ± 1.5 | 95.1 ± 1.2 |
| **24 h** | **Resazurin** | **Neutral red** | **CFDA-AM** | **24 h** | **Resazurin** | **Neutral red** | **CFDA-AM** |
| **Control** | 100.0 ± 0.4 | 100.0 ± 1.5 | 100.0 ± 0.8 | **Control** | 100.0 ± 0.8 | 100.0 ± 0.8 | 100.0 ± 1.3 |
| **PFOS 1 nM** | 99.4 ± 0.7 | 104.6 ± 1.3* | 99.0 ± 0.9 | **$CdCl_2$ 1 nM** | 99.7 ± 1.3 | 101.2 ± 1.4 | 99.6 ± 1.7 |
| **PFOS 10 nM** | 98.0 ± 0.8 | 108.5 ± 1.1**** | 95.4 ± 1.2 | **$CdCl_2$ 10 nM** | 98.5 ± 1.5 | 102.3 ± 1.8 | 102.5 ± 1.6 |
| **PFOS 100 nM** | 97.4 ± 0.6 * | 106.8 ± 0.9 ** | 94.4 ± 1.4 * | **$CdCl_2$ 100 nM** | 98.5 ± 1.5 | 101.7 ± 1.7 | 103.9 ± 2.2 |
| **PFOS 1 μM** | 98.3 ± 0.8 | 107.3 ± 1.5 *** | 92.8 ± 2.1 *** | **$CdCl_2$ 1 μM** | 97.2 ± 0.9 | 97.7 ± 1.7 | 102.7 ± 3.1 |
| **48 h** | **Resazurin** | **Neutral red** | **CFDA-AM** | **48 h** | **Resazurin** | **Neutral red** | **CFDA-AM** |
| **Control** | 100.0 ± 0.3 | 100.0 ± 0.9 | 100.0 ± 0.9 | **Control** | 100.0 ± 0.4 | 100.0 ± 0.5 | 100.0 ± 1.0 |
| **PFOS 1 nM** | 98.7 ± 0.5 | 107.5 ± 0.7 **** | 102.2 ± 1.7 | **$CdCl_2$ 1 nM** | 95.6 ± 0.8 ** | 101.2 ± 1.0 | 101.1 ± 1.1 |
| **PFOS 10 nM** | 96.8 ± 0.8 ** | 111.3 ± 1.1 **** | 98.8 ± 1.7 | **$CdCl_2$ 10 nM** | 93.0 ± 1.2 **** | 102.5 ± 1.4 | 103.1 ± 1.6 |
| **PFOS 100 nM** | 98.1 ± 0.9 | 105.9 ± 1.2 *** | 96.9 ± 2.0 | **$CdCl_2$ 100 nM** | 92.1 ± 1.2 **** | 102.7 ± 1.8 | 101.6 ± 1.8 |
| **PFOS 1 μM** | 99.3 ± 0.7 | 107.0 ± 1.5 **** | 92.8 ± 2.1 | **$CdCl_2$ 1 μM** | 96.3 ± 0.7 * | 99.7 ± 1.5 | 103.0 ± 2.5 |
| **24 h** | **Resazurin** | **Neutral red** | **CFDA-AM** | **24 h** | **Resazurin** | **Neutral red** | **CFDA-AM** |
| **Control** | 100.0 ± 0.3 | 100.0 ± 1.0 | 100.0 ± 1.4 | **Control** | 100.0 ± 0.6 | 100.0 ± 0.9 | 100.0 ± 1.0 |
| **DEHP 1 nM** | 98.8 ± 0.6 | 103.4 ± 1.5 | 100.8 ± 1.9 | **DDE 1 nM** | 98.4 ± 0.7 | 100.8 ± 1.2 | 99.4 ± 0.7 |
| **DEHP 10 nM** | 98.3 ± 0.7 | 103.9 ± 1.1 | 101.7 ± 1.7 | **DDE 10 nM** | 97.5 ± 0.8 | 101.9 ± 1.2 | 97.4 ± 0.8 |
| **DEHP 100 nM** | 97.0 ± 0.8 * | 102.5 ± 1.2 | 98.2 ± 1.5 | **DDE 100 nM** | 95.5 ± 0.7 *** | 99.7 ± 1.1 | 95.4 ± 1.2 ** |
| **DEHP 1 μM** | 98.6 ± 1.2 | 100.1 ± 2.1 | 93.5 ± 2.2* | **DDE 1 μM** | 94.5 ± 1.2 **** | 96.9 ± 1.1 | 95.7 ± 0.9 ** |
| **48 h** | **Resazurin** | **Neutral red** | **CFDA-AM** | **48 h** | **Resazurin** | **Neutral red** | **CFDA-AM** |
| **Control** | 100.0 ± 0.4 | 100.0 ± 0.8 | 100.0 ± 1.5 | **Control** | 100.0 ± 0.6 | 100.0 ± 0.4 | 100.0 ± 0.9 |
| **DEHP 1 nM** | 100.1 ± 1.1 | 102.9 ± 0.7 | 98.3 ± 2.3 | **DDE 1 nM** | 100.7 ± 0.9 | 104.8 ± 0.9 *** | 100.6 ± 1.0 |
| **DEHP 10 nM** | 99.3 ± 0.9 | 107.4 ± 1.8 * | 106.7 ± 2.8 | **DDE 10 nM** | 96.8 ± 1.6 | 106.2 ± 0.6 **** | 95.5 ± 1.6 |
| **DEHP 100 nM** | 97.5 ± 0.7 | 106.9 ± 1.6 * | 98.3 ± 1.1 | **DDE 100 nM** | 96.7 ± 1.4 | 104.6 ± 0.7 *** | 97.6 ± 1.8 |
| **DEHP 1 μM** | 101.7 ± 0.9 | 100.1 ± 2.8 | 85.9 ± 2.4 **** | **DDE 1 μM** | 96.5 ± 0.9 * | 95.3 ± 1.2 *** | 94.8 ± 1.5 * |
| **72 h** | **Resazurin** | **Neutral red** | **CFDA-AM** | **72 h** | **Resazurin** | **Neutral red** | **CFDA-AM** |
| **Control** | 100.0 ± 0.6 | 100.0 ± 1.5 | 100.0 ± 1.0 | **Control** | 100.0 ± 1.1 | 100.0 ± 1.4 | 100.0 ± 1.0 |
| **DEHP 1 nM** | 95.3 ± 1.2 * | 102.5 ± 1.9 | 91.7 ± 1.6 ** | **DDE 1 nM** | 98.9 ± 1.7 | 100.6 ± 2.5 | 100.0 ± 1.9 |
| **DEHP 10 nM** | 99.5 ± 0.8 | 104.8 ± 2.1 | 100.3 ± 2.5 | **DDE 10 nM** | 99.4 ± 1.8 | 103.6 ± 2.5 | 100.0 ± 1.8 |
| **DEHP 100 nM** | 98.5 ± 0.6 | 105.6 ± 2.1 | 98.1 ± 2.5 | **DDE 100 nM** | 100.3 ± 1.1 | 103.2 ± 2.0 | 100.7 ± 1.7 |
| **DEHP 1 μM** | 99.4 ± 1.4 | 98.3 ± 3.2 | 85.4 ± 3.0 *** | **DDE 1 μM** | 97.4 ± 0.8 | 96.7 ± 1.6 | 96.5 ± 1.5 |

**Supplemental Table 2.** Viability of EndoC-βH1 cells treated for 24, 48 or 72 h with different BPA, BPS, $CdCl_2$, PFOS, DDE or DEHP concentrations (1 nM–1 μM) as evaluated by RZ, NR and CFDA-AM assays. n= at least four independent experiments. * vs. Control; *$p < 0.05$, **$p < 0.01$, ***$p < 0.001$ and ****$p < 0.0001$, one-way ANOVA followed by Dunnet's post hoc test or Kruskal-Wallis followed by Dunn's post hoc test. All data are expressed as mean ± SEM.

| | GEN | FORWARD (5'->3') | REVERSE (5'->3') | REFSEQ (NM) |
|---|---|---|---|---|
| MIN6 | *Ins* | TTATTGTTTCAACATGGCCC | CAAAGGTGCTGCTTGACAAA | 001185083 |
| | *Pdx1* | GGCCTGGAAGAGCCCAACCG | TGTGTAAGCACCTCCTGCCCACT | 008814 |
| | *Hnf4α* | TCTGGATGACCAGGTGGCGCT | GGACACACGGCTCATCTCCGC | 008261 |
| | *MafA* | CCCGTCTTGGCCCTCCATGATT | TATCCCGCCGTGTCTGTGTGG | 194350 |
| | *Kir6.2* | CCCGTCTTGGCCCTCCATGATT | TATCCCGCCGTGTCTGTGTGG | 010602 |
| | *Sur1* | TGCCTCAGGACAAGCAACC | GACCACTGTCCTCTTGTCATC | 011510 |
| | *Glut2* | ATCGCTCCAACCACACTCAG | CTGAGGCCAGCAATCTGACTA | 031197 |
| | *Gck* | TTCAGCTTCTGGCCTCCCACAG | AAAACAGCCAGGTCTGGGCAGC | 010292 |
| | *Hprt* | GGTTAAGCAGTACAGCCCCA | TCCAACACTTCGAGAGGTCC | 013556 |
| | *Actb* | GGCTGTATTCCCCTCCATCG | CCAGTTGGTAACAATGCCATGT | 007393 |
| EndoC-βH1 | *INS* | AAGCAGATCACTGTCCTTC | ACACTAGGTAGAGAGCTTCC | 000207 |
| | *PDX1* | CCTTTCCCATGGATGAAGTC | CGAACTCCTTCTCCAGCTCTA | 000209 |
| | *HNF4α* | CGAAGGTCAAGCTATGAGGACA | ATCTGCGATGCTGGCAATCT | 178849 |
| | *MAFA* | TTCTCCTTGTACAGGTCCCG | GAGAGCGAGAAGTGCCAACT | 201589 |
| | *MAFB* | AAGCCTCTCACCCTAGGAGC | GGAAAACAGATCCTCCCCTC | 005461 |
| | *KIR6.2* | AGGTCCAAGTGACTATTGGCT | TCTGCACGATGAGGATCAGGA | 000525 |
| | *SUR1* | TCGTGAATCTGCTGTCCAAAG | CGATGGCTCGCAAGTCGAT | 001287174 |
| | *SNAP25* | TCGTGTAGTGGACGAACGG | TCTCATTGCCCATATCCAGGG | 003081 |
| | *GLUT1* | TCTGGCATCAACGCTGTCTTC | CGATACCGGAGCCAATGGT | 006516 |
| | *GCK* | TGCTACTACGAAGACCATCAGT | CCACTCGGTATTGACGCACA | 000162 |
| | *HPRT1* | GCCCTGGCGTCGTGATTAGT | AGCAAGACGTTCAGTCCTGTCCATAA | 000194 |
| | *ACTB* | CACCATTGGCAATGAGCGGTTC | AGGTCTTTGCGGATGTCCACGT | 001101 |

**Supplemental Table 3.** Primer sequences used in RT-qPCR for the study of pancreatic β-cell gene expression.
